# Untargeted metabolic analysis reveals intraspecific and organ-specific chemodiversity in *Solanum dulcamara*

**DOI:** 10.64898/2026.07.29.741464

**Authors:** Judit Valeria Mendoza-Servín, Abigail Moreno-Pedraza, Paula Carolina Pires Bueno, Nicole M. van Dam

## Abstract

**Background and Aims:** The genus *Solanum* including the wild species *S. dulcamara*, is rich in specialized metabolites such as steroidal glycoalkaloids (SGAs). Yet, much of this chemical diversity remains poorly characterized. This study aims to provide a comprehensive assessment of intra-specific chemodiversity in *S. dulcamara*. Using a dataset generated from 42 globally distributed accessions, we tested whether metabolic profiles differ among plant organs. We postulated that metabolic richness and abundance vary across accessions. Additionally, we hypothesized that differences in geographic origin or altitude affect SGA chemodiversity.

**Methods:** An untargeted metabolomic approach was applied to leaf, flower and root samples of 42 *S. dulcamara* accessions. Plants were grown in the greenhouse, and the extracted metabolites were analyzed using UHPLC-HRMS/MS in positive and negative ionization modes. Data processing and metabolite annotation were performed with a tailored bioinformatics workflow. Multivariate analyses were performed to evaluate chemical variation across organs and accessions.

**Key Results:** Our analyses revealed both organ and accession-specific metabolic diversity. Principal component analysis and clustering analyses revealed metabolic differentiation between leaves, flowers and roots. Leaves showed the highest metabolite richness and abundance, while roots showed the lowest. Alkaloids, especially SGAs, dominated positive mode profiles in roots, whereas shikimates and phenylpropanoids were prominent in negative mode profiles. Based on the leaf and flower SGAs profiles, four chemotypes were identified. Analyses of flavonoid and cinnamic acid derivatives, however, did not reveal chemotypes. Feature-based molecular network analyses confirmed that metabolite clusters are associated with plant organs, but not with altitude or geographic origin of the accessions.

**Conclusions:** The intraspecific chemodiversity within *S. dulcamara* is mainly driven by organ and accession-specific metabolic differences. We identified four SGA leaf and flower chemotypes, suggesting possible functional and ecological roles of this aboveground chemodiversity. These insights may contribute to applied research in plant resistance breeding and crop production.

## INTRODUCTION

Plants produce an extraordinary number of organic compounds through their metabolism. Some, known as primary metabolites, are essential for survival and growth; others, the so-called secondary or specialized metabolites, help plants adapt to environmental conditions (Erb & Kliebenstein, 2020). These phytochemicals mediate interactions with herbivores, pathogens, mutualists, and competitors. Their composition can vary widely among individuals, tissue types, and developmental stages (Anaia et al., 2024; van Dam, 2009b). Yet for many of these compounds, we lack even the most basic information, because a vast number of molecules have not been isolated nor structurally described. This means that their functions remain unknown, and we still do not understand how their production and concentrations are affected due to natural selection pressures and changing environmental conditions (Petrén et al., 2024). Addressing these questions requires studying plant groups with high chemical diversity and ecological variability.

The genus *Solanum* is one of 99 genera within the Solanaceae family and represents one of the largest and most taxonomically complex groups of angiosperms. Approximately 2,000 species have been described worldwide, spanning all continents and encompassing a wide range of climatic conditions. Several globally important crops belong to this genus, including *S. lycopersicum* (tomato), *S. tuberosum* (potato), and *S. melongena* (eggplant) (Kaunda & Zhang, 2019; Poczai et al., 2022). Beyond their agricultural importance, members of the Solanaceae family are recognized for their remarkable diversity of specialized metabolites, which drives their well-established use in traditional medicine (Kaunda & Zhang, 2019). The chemical diversity of *Solanum* makes this genus an important system for studying plant secondary metabolism and its resulting ecological interactions and functions.

Within this context, the wild nightshade *Solanum dulcamara* L. (bittersweet nightshade) is a particularly useful model for exploring intra-plant chemical diversity. Native to Europe and western Asia, *S. dulcamara* occupies a broad range of habitats, including wetlands, riparian zones, forests, gardens, roadsides, and even relatively dry environments (Savinykh & Konovalova, 2019). Chemically, *S. dulcamara* is characterized by the production of steroidal glycosides (SGs), a group of bioactive and toxic specialized metabolites that include steroidal glycoalkaloids (SGAs) and steroidal saponin glycosides (SSGs), both widely distributed within Solanaceae (Anaia et al., 2024; Chiocchio et al., 2023). Steroidal glycosides consist of a hydrophobic steroidal backbone linked to one or more sugar moieties. These toxic compounds are synthesized from cholesterol through sequential hydroxylation, oxidation, transamination, and glycosylation reactions, many catalyzed by enzymes encoded in the GAME (GlycoAlkaloid Metabolism) gene clusters (Itkin et al., 2013). Functionally, SGAs interact with membrane sterols and can disrupt membrane integrity by depolarizing membrane potential and interfering with active ion transport (Blankemeyer et al., 1997; Roddick & Drysdale, 1984) thereby contributing to plant defense. Previous studies showed that *S. dulcamara* accessions differ in their SGs profiles (Calf et al., 2018; Willuhn, 1966). Using seed batches collected in different Dutch *S. dulcamara* populations, Calf et al., (2018) identified distinct SGA leaf chemotypes, one even lacking common SGAs. These (heritable) chemotypes differed in their resistance to herbivores, in particular slugs and specialist flea beetles (Calf et al., 2018, 2019) A more detailed study on two chemotypes containing mostly saturated or unsaturated SGAs showed that chemotype formation was linked to differential expression of specific GAME genes (Anaia et al., 2024). While steroidal glycosides represent a well-characterized class of defense compounds in *Solanum*, they constitute only a fraction of the overall chemical diversity present in plants. To achieve a more comprehensive understanding of the evolutionary drivers and ecological consequences of plant chemodiversity, broader metabolic investigations are required, and advances in analytical technologies have enabled chemical ecology to move in this direction.

Untargeted metabolomics is a global, comprehensive approach that aims to analyze and quantify a broad range of low-molecular-weight molecules in a sample; however, physicochemical properties, technological development, and methodological pipelines limit the number of detected compounds (Souza & Patti, 2021). This approach makes it possible to detect and characterize a wide range of phytochemicals in complex biological matrices, even when many compounds co-occur in a single sample (Di Minno et al., 2021). Subsequent biological interpretation depends on metabolite identification. This step involves computationally matching experimental mass spectra against a reference database to generate putative identities (Schrimpe-Rutledge et al., 2016). Continuous improvements in data processing workflows and expanding spectral resources are progressively enhancing our ability to interpret metabolomic data, thereby strengthening its application in ecological and evolutionary studies (Uthe et al., 2021; Zulfiqar et al., 2023).

In this study, we applied untargeted metabolomics to analyze intraspecific chemical diversity involving the extraction, detection and analysis of low molecular weight primary and secondary metabolites in *S. dulcamara* across 42 accessions from multiple geographic locations. We grew all the plants under similar conditions in a greenhouse and sampled roots, leaves and flowers. The samples were analyzed using Ultra High-Performance Liquid Chromatography coupled to tandem High Resolution Mass Spectrometry (UHPLC-HRMS/MS) in both positive and negative modes. Using this comprehensive *S. dulcamara* metabolic data set, we tested three hypotheses: (1) organs differ in their metabolic profiles, reflecting organ-specific metabolic specialization; (2) steroidal glycoalkaloids (SGAs) are a major driver of chemodiversity, enabling to assign accessions to the previously described *S. dulcamara* leaf steroidal glycoside chemotypes; and (3) geographic origins or altitude consistently affect the metabolic profiles of the different accessions.

## MATERIALS AND METHODS

### Chemicals and reagents

LC-MS grade acetonitrile and water were purchased from Merck (Darmstadt, DE), and LC-MS grade formic acid was purchased from Tokyo Chemical Industry Co. (Tokyo, JP). The internal standard ethyl gallate (C_9_H_10_O_5_, MW 198.17 g/mol, ≥96.0%), was purchased from Sigma-Aldrich, Co. (St. Louis, MO, USA).

### Plant material

A total of 42 accessions of *S. dulcamara* were obtained from different sources across eleven different geographic locations. For unnamed accessions, we assigned acronyms based on geographic origin - country followed by city or location (e.g., GBS: Germany, Berlin, Spree River) - as described in Supplementary Information S1. The accessions obtained as seeds were sown in 11.5 x 10 x 8 cm plastic boxes containing a 2 cm layer of glass beads (5 mm Ø) almost covered with tap water. The seeds were cold stratified in the dark at 4 °C for two weeks (Chiocchio et al., 2023). After stratification, boxes were transferred to a greenhouse cabin (16 h light/8 h dark photoperiod at a 17-25 °C). When seedlings had developed two cotyledons, they were transplanted into pots filled with 1:1 (v/v) soil:sand mixture and fertilized with 4 g.L⁻¹ of slow-release fertilizer (Osmoeote Extract Standard, Everris, International B.v.). Each accession was represented by a single individual plant, which was clonally propagated by preparing six stem cuttings (∼15 cm long). Cuttings were placed in 50 mL Falcon tubes containing 15 mL tap water, where they developed adventitious roots and new leaves. The rooted cuttings were then transplanted into pots. Once plants reached ∼40 cm height, wooden sticks were used for support.

### Sample preparation

For metabolite extraction, three clonal plants were sampled for vegetative organs (leaves and roots), and the remaining three were sampled for reproductive organs (flowers). Roots were harvested after gently washing off adhering soil. Four fully expanded leaves at a defined ontogenetic position (leaves 7 to 10, counted from the apex) were excised with disinfected scissors. Between 8 and 15 fully open flowers were harvested. All samples were immediately flash-frozen and kept at -70 °C until freeze-drying.

Freeze-dried samples were ground in a ball mill (Mixer Mill MM 400; Retsch) with two metal beads (4 mm Ø, 30 Hz, 1.3 min). For extraction (including primary and secondary metabolites), 10 mg of each sample was weighed into a 2 mL Safe-Lock tube (Eppendorf, Germany). Samples were extracted with 1 mL of extraction solution consisting of 25% water and 75% methanol (v/v) spiked with 25 ppm of ethyl gallate used as the internal standard (IS). Two metal beads (4 mm Ø) and samples were shaken at 30 Hz for 3 minutes in the ball mill, followed by sonication for 15 min in an ultrasonic bath (Sonorex Super Rk). Sample extracts were centrifuged at 3000 rpm for 2 min at room temperature, supernatants were filtered through a 0.2 μm polytetrafluoroethylene (PTFE) membrane and transferred to new 1.5 mL glass vials fitted with 200 µL inserts. Quality control (QC) was prepared with a mix of all the samples in equal parts.

### UHPLC-HRMS/MS data acquisition and processing for untargeted metabolomics analysis

For the untargeted metabolomics analysis, Ultra High-Performance Liquid Chromatography (UHPLC) coupled to tandem High Resolution Mass Spectrometry (HRMS/MS) was employed. A volume of 0.5 μL of each extract was injected into an Agilent 1290 Infinity UHPLC system coupled to an Agilent 6546 Q-TOF mass analyzer (Agilent Technologies). Chromatographic separation was performed on a C18 analytical column (InfinityLab Poroshell 120 EC-C18, 2.1 x 100 mm, 1.9 μm, 120 Å, Agilent Technologies) protected by a guard column of the same stationary phase. The column temperature was set to 35 °C and the flow rate at 0.42 mL/min. The mobile phase consisted of (A) water containing 0.05% (v/v) formic acid and (B) acetonitrile containing 0.05% (v/v) formic acid, following the gradient: from 0 to 1 min at 2% of B, from 1 to 8 min from 2 to 20% of B, from 8 to 8.5 min from 20 to 40% of B, from 8.5 to 12 min from 40 to 42% of B, from 12 to 20 min from 42 to 45% of B, from 20 to 22 min from 45 to 50% of B, from 22 to 22.5 min from 50 to 98% of B, from 22.5 to 25 min keeping at 98% of B, from 25 to 26 min from 98 to 2% of B and from 26 to until 29 min reconditioning at 2% of B.

The mass spectrometric data were acquired in positive and negative ionization modes using an Agilent Jet Stream ESI source operating in Data Dependent Acquisition (DDA) mode. For both modes, the capillary voltage was ±4 kV, drying gas at 220 °C (11 L.min⁻¹), nebulizer at 40 psi, and sheath gas at 375 °C at a gas flow of 12 L.min⁻¹. Specific source parameters included: nozzle voltage at 0 V, Fragmentor at 100, Skimmer at 65, and Octopole RFPeak at 750. The full scan range was set from 60 to 1500 m/z for both MS1 and MS2. During DDA, a maximum of 2 precursor ions (intensity threshold > 200) were isolated and fragmented using collision-induced dissociation (CID) energies of 5, 10, 20, 30, and 40 eV at 5 spec/s for positive mode and 20 eV for negative mode. Accuracy was maintained by continuous recalibration using reference ions (121.050873 and 922.009798 Da in positive ionization mode and 112.985587 and 966.000725 in negative ionization mode).

The raw mass spectrometry files (.d) were converted to an open format (.mzML) using MSConvert 3.0 from ProteoWizard (Chambers et al., 2012) with peak picking enabled. Data were imported into the open-source software MZmine 4.8.30 (Schmid et al., 2023) and parameters were adjusted to the instrument specifications and ESI polarity. Features with low reproducibility in the QC injections were filtered using a maximum relative standard deviation (RSD) threshold of 40%. Solvent blanks (MeOH) were used for blank subtraction. Putative compound annotations were obtained by automated database (DB) matching in MZmine based on accurate mass and MS/MS spectral similarity, additional evidence was obtained through spectral library searches and a home-made database containing characteristic metabolites reported for *Solanaceae* species, including compounds registered in FoodDB (https://foodb.ca) with a confidence level of 2 based on the Metabolomics Standards Initiative (MSI) (Sumner et al., 2007). MS/MS spectra were then exported in Mascot Generic Format (.mgf) and further processed in SIRIUS 6.0 (https://bio.informatik.uni-jena.de/software/sirius/) to support putative structure identification and compound class prediction. Finally, all features of interest were manually inspected by comparing their MS/MS fragmentation patterns with spectral libraries, SIRIUS predictions, and published fragmentation data. When the experimental fragmentation patterns supported a specific metabolite assignment, the initial automated annotation was replaced by a manually curated putative metabolite annotation (MSI Level 2). No compounds were assigned to Level 1 since no reference standards were used. The final outputs were combined and organized as feature-intensity matrices (relative areas) for statistical analyses.

### Data analyses

Metaboanalyst 6.0 (https://www.metaboanalyst.ca/) (Pang et al., 2024), and RStudio (Posit team, 2025) were used for data analysis and statistics. Feature lists from MZmine were normalized to sample weight and to the IS, followed by log_2_ transformation and Pareto scaling, except when explicitly mentioned otherwise. Samples were analyzed separately for datasets generated in positive (ESI+) and negative (ESI−) ionization modes.

Principal Component Analysis (PCA) was performed using MetaboAnalyst 6.0. Venn Diagrams were created using the *VennDiagram* R package (H. Chen & Boutros, 2011). Mascot generic files were imported into SIRIUS 6.3.0 (Dührkop et al., 2019) for in-silico annotation, used to assign features into chemical classes by using the Natural Product Classifier (NPC). For doughnut and sunburst plot analyses, data were used without log₂ transformation or Pareto scaling. These plots were generated using *tidyverse* and *plotly* R package (Sievert, 2020; Wickham et al., 2019). Individual features from the same organ and ionization mode, biological replicates were averaged and filtered with a confidence probability > 0.8 at the pathway, superclass and class levels (NPC) (Supplementary Information S2 and S3).

To generate hierarchical clustering, mean peak intensities were calculated for the three biological replicates of each accession and organ type. To compare the hierarchical clustering patterns between the different organs, a tanglegram was constructed using the *dendextend* R package (Galili, 2015). Before averaging, hierarchical clustering utilizing Euclidean distance and the Ward clustering algorithm was performed to generate dendrograms, ensuring the high reproducibility of replicates within each sample group.

Richness was calculated as the number of unique features detected per accession. Differences in richness were then modeled using generalized linear models with a negative binomial distribution to account for count structure and overdispersion, including accession, organ type, and their interaction as fixed effects. Metabolite abundance was calculated as the sum of all feature peak intensities per accession. Total metabolite abundance was analyzed using linear models. Model selection was performed using Akaike Information Criterion (AIC), and final model significance was assessed using Type III ANOVA. Post-hoc pairwise comparisons were conducted using estimated marginal means (emmeans). Model assumptions were evaluated using diagnostic plots and the DHARMa package (Hartig, 2022).

To visualize metabolite distribution across accessions, heatmaps were generated for each organ type and chemical class using the *ComplexHeatmap* R package (Gu et al., 2016). Raw data were normalized per organ separately and then Z-score normalized by row to highlight relative abundance differences between samples. Features with zero standard deviation or entirely missing values after scaling were removed to ensure numerical stability. Any remaining missing values or infinite results were imputed with 0 to facilitate Euclidean distance calculations. Three major chemical groups were prioritized for analysis based on their relative abundance. For compounds detected in ESI+ mode, SGAs were manually annotated by integrating retention time (RT), high-resolution mass spectrometry accurate mass, and specific in-source fragmentation patterns. Flavonoids and phenylpropanoids detected in ESI-were identified using the NPC class level. Only annotations with confidence probability greater than 0.8 were retained, and putative names were assigned based on spectral matching within the MZmine framework.

A feature-based molecular networking (FBMN) workflow was performed within the Global Natural Products Social Molecular Networking (GNPS2, http://gnps2.org/homepage, release 2025.12.10) for data organization and visualization according to the plant organs and geographical information. Cytoscape version 3.10.3 was used for graphical presentation.

## RESULTS

### Chemical diversity of S. dulcamara varies across roots, leaves and flowers

Untargeted metabolomics was used to examine the chemical profile of roots, leaves and flowers of 42 *S. dulcamara* accessions by UHPLC-HRMS/MS, using electrospray ionization (ESI). To achieve broader metabolite coverage, both positive (ESI+) and negative (ESI-) ionization modes were applied. Following extraction, acquisition, and data processing, the resulting datasets from the three organs yielded a total of 3,690 features in positive mode and 2,373 features in negative mode.

The resulting feature intensity matrices representing relative peak areas from each ionization mode were analyzed using PCA. For data acquired in positive ionization mode, the first two principal components in the score plot explained 43.6% of the total variance (PC1 = 30.2% and PC2 = 13.4%). The separation among organs was mainly driven by PC1, with roots forming a well-separated cluster on the right side of this component. Additionally, within the larger leaf and flower clusters, we observed two sub-clusters separating along PC2 (Figure 1A, ESI+). For features detected in negative ionization mode, the first two principal components explained 54.1% of the total variance (PC1 = 41.5% and PC2 = 12.6%). The PCA revealed well defined clusters along both PC1 and PC2, allowing a clear separation among organs (Figure 1A, ESI-).

**Figure 1.**
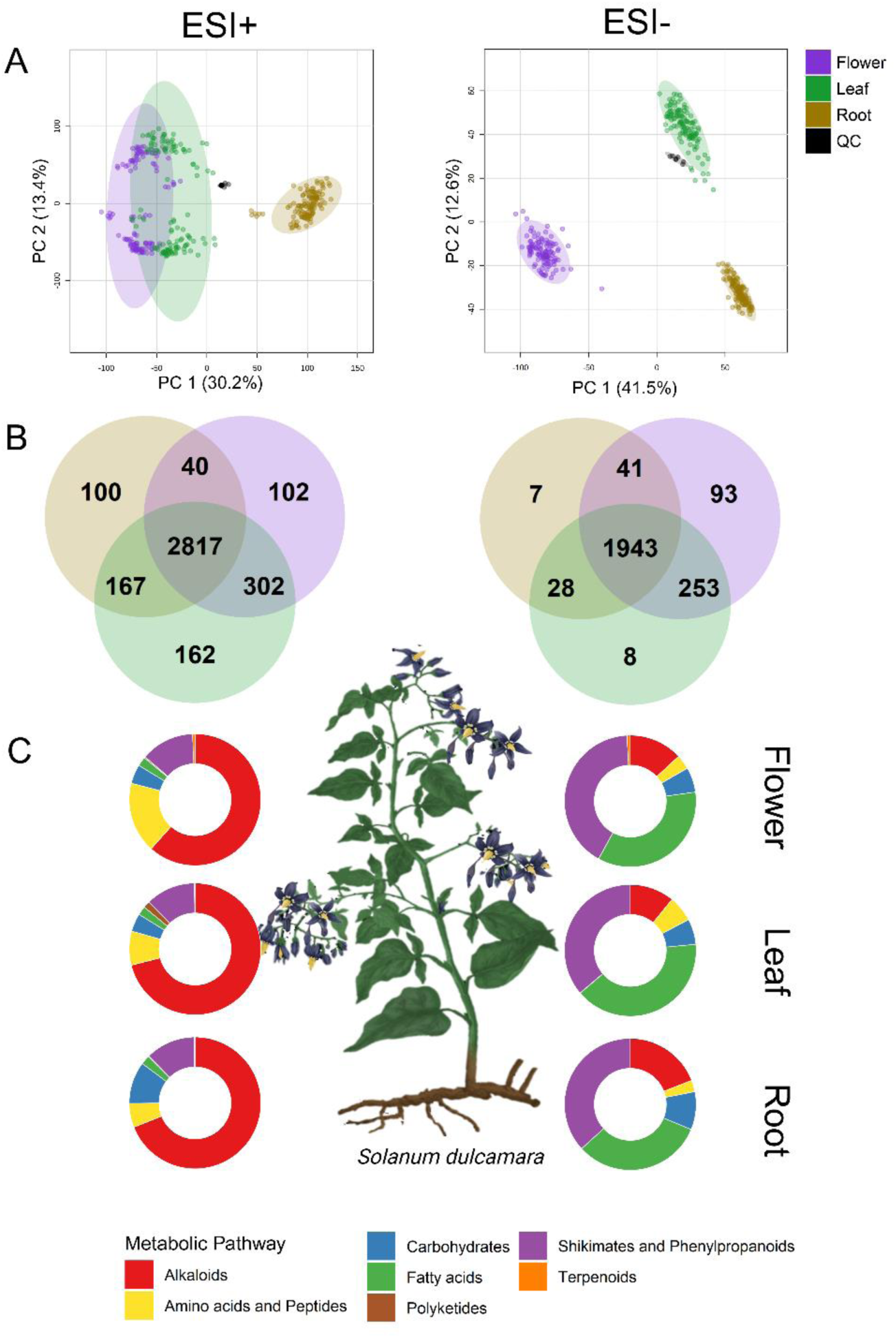
Organ-specific metabolome profiling across positive (ESI+) and negative (ESI-) ionization modes in *Solanum dulcamara*. (A) Principal Component Analysis (PCA) plots of features detected in ESI+ (right) and ESI- (left) showing a clear separation between flower (purple), leaf (green), and root (light brown) samples; the QC cluster indicates good analytical reproducibility (black color). Explained variances in PC1 and PC2 are shown in the axes. (B) Venn diagrams summarizing shared and organ-enriched metabolite features between flower, leaf, and root in ESI+ (left) and ESI− (right). (C) Donut charts showing the relative distribution of annotated metabolic pathways (alkaloids, amino acids/peptides, carbohydrates, fatty acids, polyketides, shikimates/phenylpropanoids, and terpenoids) in flower, leaf, and root in ESI+ (left column) and ESI- (right column).

Most of the detected features were shared among the three organs, indicating a large common metabolic core (Figure 1B). In positive mode (Figure 1B, ESI+), 76.3% of the detected features were shared among the three organs. The overlap was greater between flowers and leaves, with 302 features shared; roots were the chemically most distinct organ, sharing 167 features with leaves and only 40 with flowers. In addition, organ-specific features were observed in leaves (162), flowers (102), and roots (100). In negative ionization mode (Figure 1B, ESI-), 81.8% of the features were shared among all three organs. Flowers exhibited the highest number of unique features (93), whereas roots and leaves showed fewer organ-specific features (7 and 8, respectively). The overlap among flowers and leaves (253) was again the most pronounced, followed by roots and flowers (41) and roots and leaves (28).

### Alkaloids, phenylpropanoids and fatty acids dominate the metabolic profile of S. dulcamara

Across both ionization modes, features were classified into seven major biosynthetic pathways. From the 3,690 features detected in ESI+, SIRIUS annotated 732, of which 249 were assigned with a confidence probability > 0.8 at the pathway, superclass and class (NPC). In ESI-, 2,373 features were detected; SIRIUS annotated 987, including 239 with confidence > 0.8 at the pathway, superclass and class (MSI Level 3).

Pathway enrichment analysis indicated that in the data acquired in ESI+, alkaloid-related metabolites represented the predominant chemical class across all three organs, accounting for the major proportion of the total signal intensity (Figure 1C). Within them, polyamines represented the most abundant class in leaves and flowers, followed by steroidal alkaloids, which dominated over all other groups (Figure 2, left side). In roots, the steroidal alkaloids predominated over all other compound groups (Figure 2, left side). After alkaloids, shikimates and phenylpropanoid derivatives represented the second most abundant pathway group in leaf and root, represented mainly by the classes of gallotannins and cinnamic acid derivatives. In flowers, they came just after amino acids and peptides.

**Figure 2.**
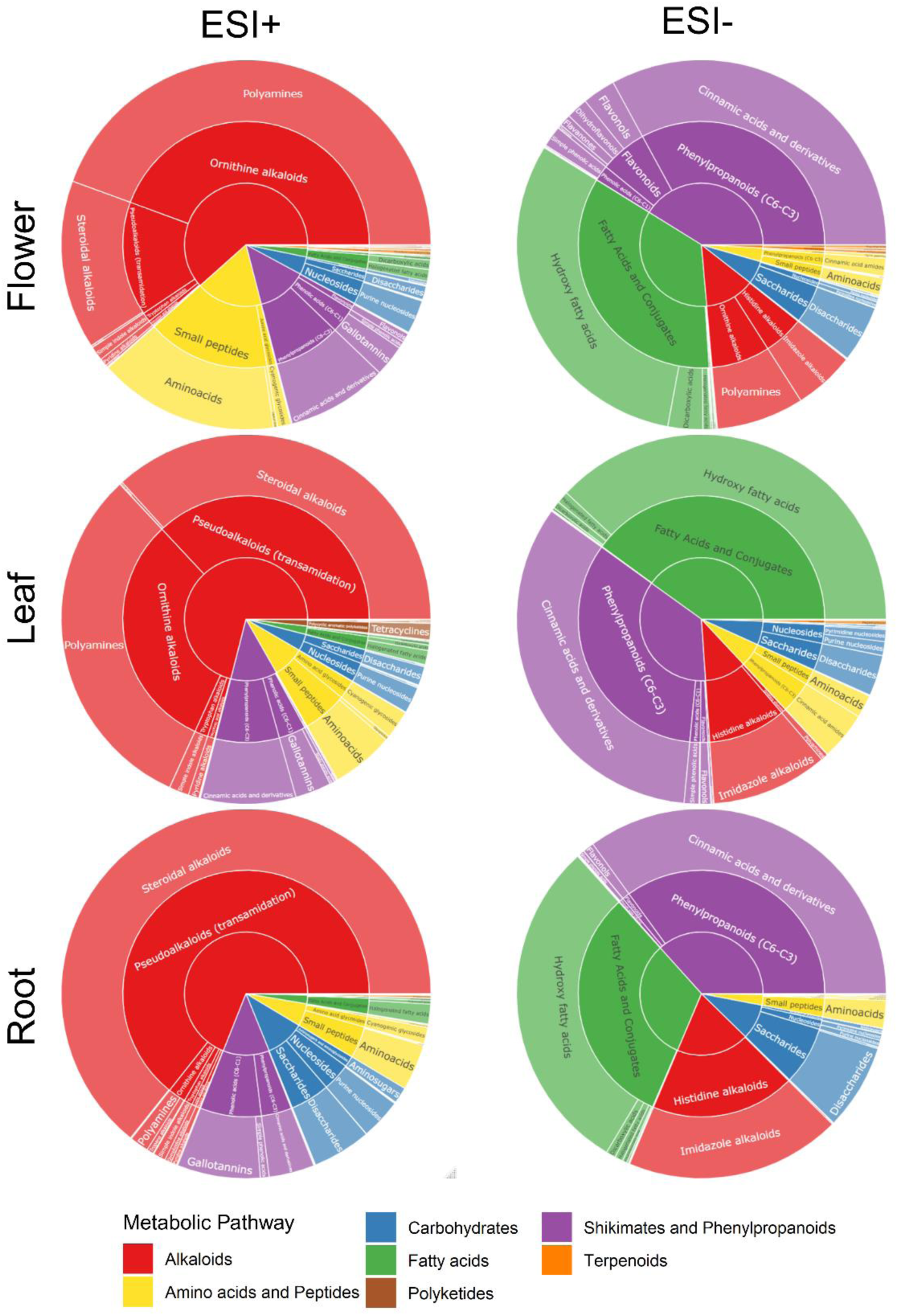
Chemical class composition of annotated metabolites across organs and ionization modes in *Solanum dulcamara*. Sunburst plots summarize the hierarchical distribution of annotated metabolite classes in flower, leaf, and root extracts acquired in ESI+ (left) and ESI− (right). Inner rings represent major pathway-level classes (color-coded), and outer rings show subclass annotations; segment size reflects the relative abundance/feature contribution within each organ and mode.

When data were acquired in ESI-, a different picture on the metabolic composition of the three organs arose. Flowers and roots showed higher average intensities of shikimates and phenylpropanoid derivatives, while fatty acids were slightly more abundant in leaves (Figure 1C, right side). Alkaloids ranked third in abundance in ESI-profiles. Among them, imidazole alkaloids constituted the main class detected in the leaves and roots, while polyamines were mostly detected in the flowers (Figure 2, right side). In flowers, flavonoids were also prominently present. In contrast, terpenoids and polyketides were the least abundant chemical classes in metabolic profiles (Figure 2). Together, these findings indicate that in both ionization modes we found substantial and complementary chemical diversity among organs, either with a focus on alkaloids (positive mode) or shikimates, phenylpropanoids and fatty acids (negative mode). Whereas all organs share most of their metabolites in a core metabolome, we also found organ-specific metabolites, particularly in the positive mode.

### Organ-specific differences in chemotype clustering patterns

To further assess similarities and differences in metabolic profiles across organs and accessions, we applied Euclidean distance and the Ward clustering algorithm analysis. For the data acquired in ESI+, the leaf and flower metabolomes of the 42 accessions were partitioned into two major clusters (Supplementary Information, Figures 1, 2). This agrees with the grouping patterns observed in the PCA using the ESI+ results. However, for root samples analyzed in ESI+ mode, a third distinct cluster emerged, comprising the samples of accessions ZD11 and ZD04 (Supplementary Information Figure 1, roots). These two accessions also formed independent clades within one of the two main clusters in both leaf and flower datasets (Supplementary Information, Figure 1, leaves and flowers). In ESI-, this additional cluster was less clearly separated, particularly in flowers, indicating a reduced discrimination among these accessions under negative ionization conditions (Supplementary Information, Figures 2).

A tanglegram analysis was conducted to evaluate the consistency of clustering patterns between organs. Since leaves and flowers had more overlaps in their metabolomes (Figure 1 A, B), we focused on the comparison of roots vs. leaves and leaves vs. flowers. In ESI+ mode, the comparison between leaves and flowers exhibited a high degree of concordance, as indicated by a greater prevalence of parallel and minimally intersecting linkage lines (Figure 3). This reflects a relatively similar positioning of accessions across both dendrograms. In contrast, the roots vs. leaves tanglegrams displayed more crossing links, suggesting substantial differences in how samples of these organs were clustered in dendrogram (Figure 3). A similar overall trend was observed in ESI-mode; however, these patterns were less pronounced. Nevertheless, leaves vs. flowers tanglegrams still demonstrated relatively greater agreement compared to roots vs. leaves tanglegrams; the linkage structure appeared more irregular, with a higher number of lines crossing (Figure 3).

**Figure 3.**
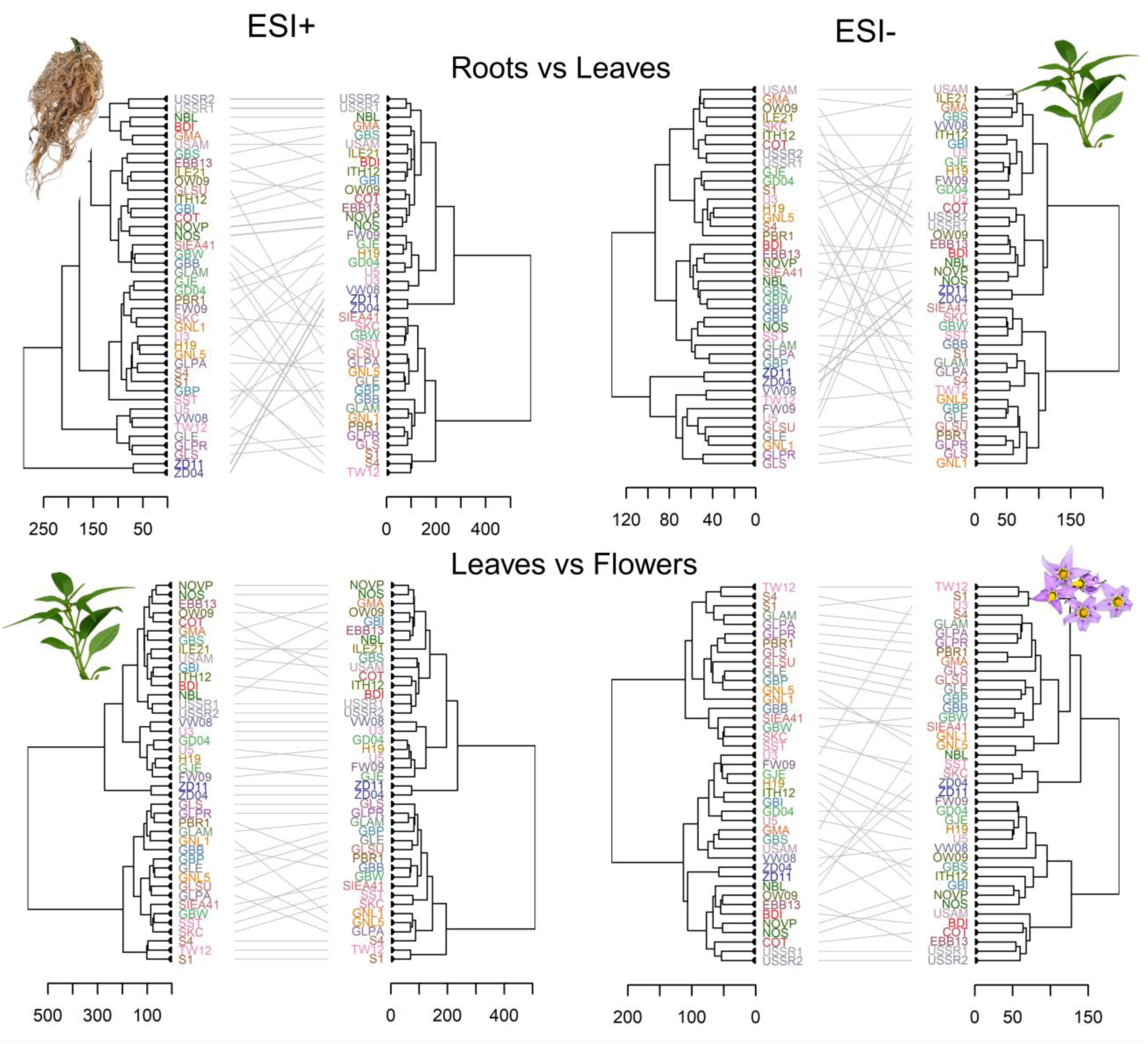
Tanglegrams compare hierarchical clustering across organs *Solanum dulcamara*. Paired dendrograms display the clustering of samples based on extracts acquired in ESI+ (left) and ESI− (right) metabolomics data for two organ comparisons: Roots vs Leaves (top panels) and Leaves vs Flowers (bottom panels). Grey links illustrate agreement and discordance in clustering structure across the two analyses.

### Chemodiversity differs across organs and accessions

When analyzing the data acquired in ESI+, we found that the chemical richness differed significantly among accessions (binomial GLM; χ² = 632.16, df = 41, p < 0.001) and organs (χ² = 27.39, df = 2, p < 0.001), with a strong accession × organ interaction (χ² = 595.98, df = 82, p < 0.001; Figure 4A; Supplementary Information S4 and S5). The metabolites’ abundance differed significantly among accessions (F₄₀,₂₄₄ = 4.34, p < 0.001), among organs (Type III ANOVA F-test from linear model; F₂,₂₄₄ = 3.59, p = 0.029), and showed a significant accession × organ interaction (F₈₀,₂₄₄ = 3.27, p < 0.001; Figure 4B). In positive ionization mode, while a strong accession x organ interaction limited the statistical resolution of individual pairwise comparisons due to conservative post-hoc corrections, Figure 4A captures a distinct, cross-accession trajectory where leaves exhibit a consistent biological trend toward higher metabolic richness. This overarching trend remains evident across most accessions despite the lack of localized pairwise significance, a phenomenon well-documented under stringent multi-testing constraints (Ho et al., 2019).

**Figure 4.**
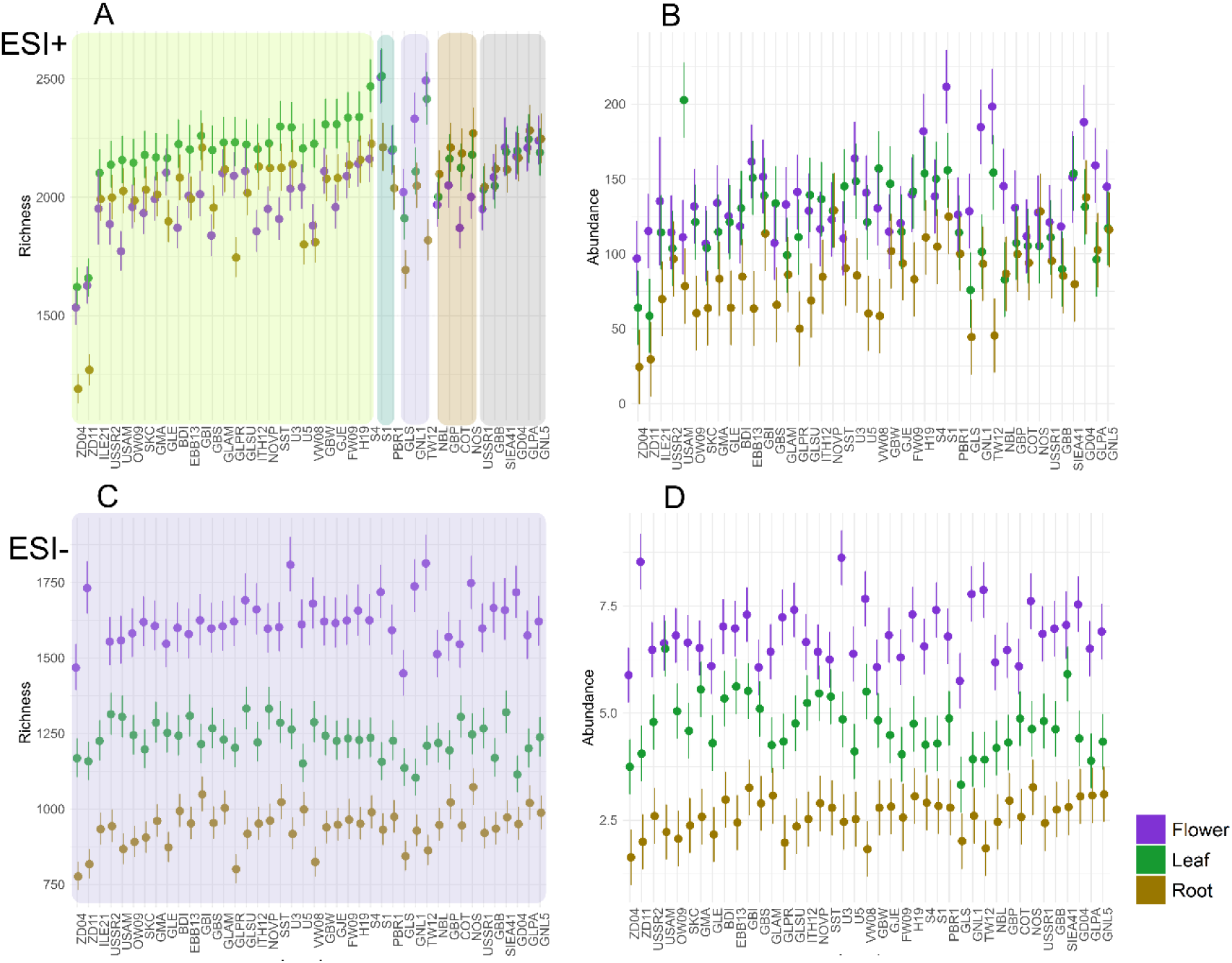
Metabolite richness and abundance in *Solanum dulcamara* across accessions and organs in positive and negative ion modes. Richness was calculated as the number of unique features detected per accession, which indicates the chemical diversity of the accession or organ. Metabolite abundance was calculated as the sum of feature peak intensities per accession, which provides information on the total concentration of each feature in the sample. Dot plots (mean ± SD) show feature richness (A) and overall abundance (B) for flower (purple), leaf (green), and root (brown) samples profiled by LC–MS in ESI+ (top row) and ESI− (bottom row). In panel A, background colors group accessions according to their richness patterns: green, purple, and brown indicate that leaves, flowers, or roots, respectively, have the highest richness; blue indicates similar richness across two or more organs; and grey indicates little differentiation in richness among organs. Accessions are ordered along the x-axis.

In ESI− mode, chemical richness also varied significantly among accessions (binomial GLM; χ² = 138.09, df = 40, p < 0.001) and organs (χ² = 157.76, df = 2, p < 0.001), with a strong accession × organ interaction (χ² = 328.05, df = 80, p < 0.001). Flowers exhibited the highest metabolic richness across all accessions, followed by leaves and roots (Figure 4C). Similar results were observed for metabolite abundance, with significant effects of accession (ANOVA; F₄₀,₂₄₄ = 4.26, p < 0.001), organ (F₂,₂₄₄ = 39.52, p < 0.001), and their interaction (F₈₀,₂₄₄ = 3.15, p < 0.001; Figure 4D).

Together, these results show that both accession and plant organs jointly determine metabolic diversity and abundance, with more consistent organ-specific patterns observed in ESI− than in ESI**+.**

### Steroidal glycoalkaloids chemotypes are not mirrored by other metabolite classes

Finally, we zoomed into the most abundant secondary metabolite classes contributing to accession- and organ-specific metabolic differentiation. We generated heatmaps to visualize the relative distribution of SGAs, flavonoids, and cinnamic acid derivatives in flowers, leaves, and roots (Supplementary Information S6-S8, Supplementary Information Figures 3-11).

Based on the heatmap of the leaf samples, the SGAs are grouped into three clusters (Figure 5, left upper heatmap). Cluster I (black) contains the SGAs with saturated (S) aglycones, including soladulcidine or tomatidine-lycotetrose, soladulcidine or tomatidine-chacotriose, their isomers, and one fragmented aglycone (possibly generated by in source fragmentation). Cluster II (dark grey) contains SGAs with unsaturated (U) aglycones, such as tomatidenol-solatriose, solasodine-chacotriose, their isomers and two fragmented aglycones. Cluster III (light grey) contains isomers of both U and S SGA aglycons present in Clusters I and II (Figure 5, left).

**Figure 5.**
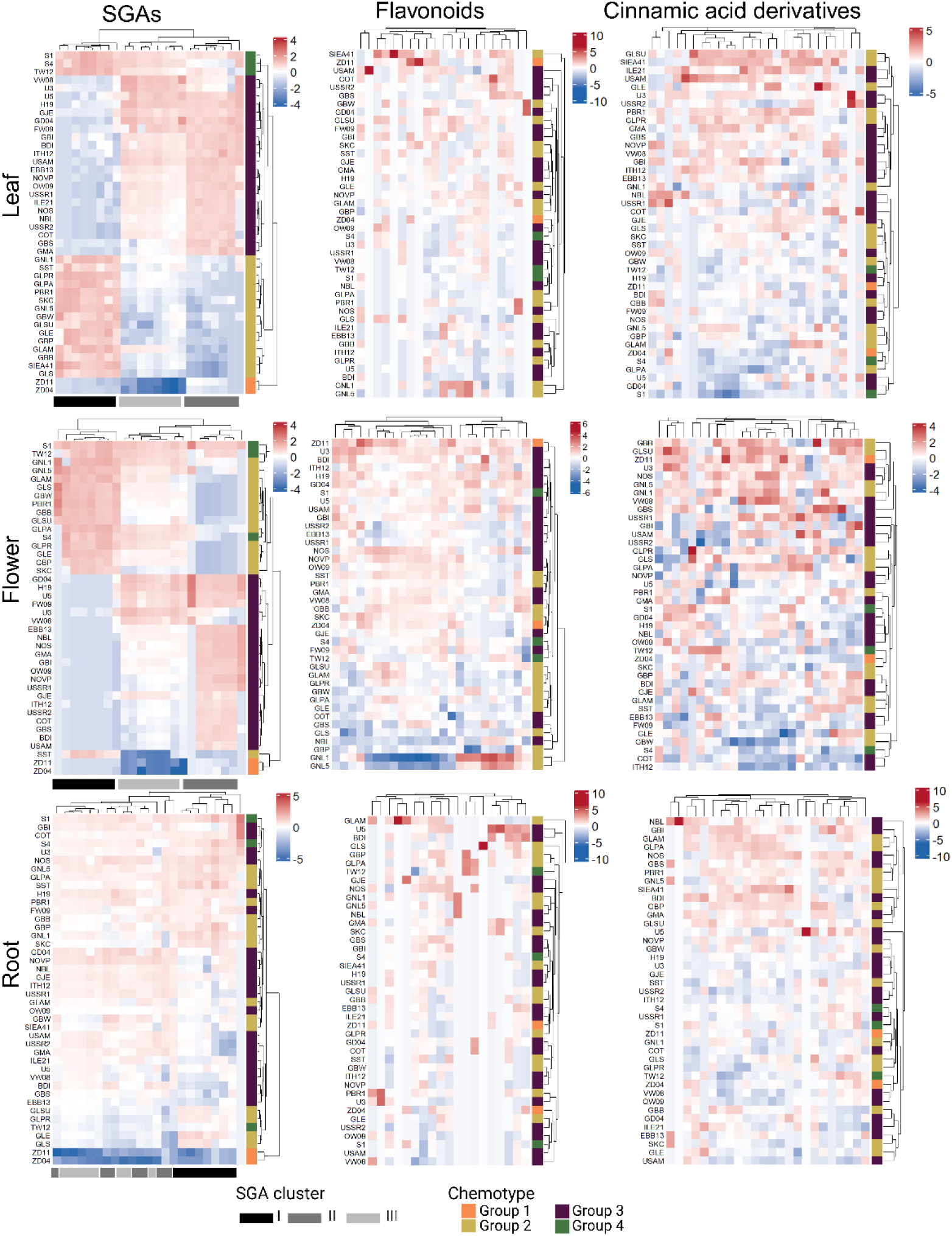
Organ-wise clustering of major specialized metabolite classes in *Solanum dulcamara.* Hierarchically clustered heatmaps show standardized metabolite levels (row z-scores; blue = lower, red = higher) for steroidal glycoalkaloids (SGAs), flavonoids, and cinnamic acid derivatives in flower, leaf, and root samples. Rows represent accessions and columns represent metabolites; dendrograms indicate similarity-based clustering, grey bars on the bottom of SGA heatmaps represent clusters based on aglycon saturation level (Cluster I: saturated, Cluster II: unsaturated, Cluster III: both saturated and unsaturated). Colored side bars denote chemotypes (Group 1–4) based on the clustering in the SGA leaf heatmap. These colors are maintained for the accessions in the other heatmaps.

Based on the leaf SGA heat maps, we also identified four chemotype groups (Figure 5, left, leaf heatmap). These were marked with separate colors and tracked in the other heatmaps. Group 1 (orange), composed of ZD04 and ZD11, showed low or undetectable SGA levels. Group 2 (yellow) shows a high abundance of SGAs from Cluster I, but low abundances of SGAs from Clusters II and III. Group 3 (brown), shows the opposite pattern, with high levels of SGA clusters II and III, but less of SGA cluster I. This group includes accessions U3 and U5, previously classified within the unsaturated (U) SGA leaf chemotype (Anaia et al., 2024). Group 4 (green) combines accessions with a relatively high abundance of all three SGA clusters. This group includes accessions S1 and S4 as well as TW12, which was previously reported to contain saturated and unsaturated SGAs in equal proportions (Calf et al., 2018).

Using the coloring of the leaf SGA chemotype groups, we found similar clusters and thus chemotypes in the flowers (Figure 5). In roots, this clustering pattern was not preserved, with only the ZD group retaining its coherence. Across roots, the abundances of the different SGAs were comparatively homogenous among accessions.

For flavonoids and cinnamic acid derivatives, no clear clustering by accession or compound class was evident (Figure 5, middle and right). However, the white flowers from accessions GNL1 and GNL5, showed a higher relative abundance of several flavonoid compounds, including peonidin glucoside, flavanones, delphinidin glucoside, quercetin glucoside, and naringenin glucoside, compared to the purple-colored flowers of the other accessions (Figure 5).

### Geographical distribution

To study whether geographic origins or altitude consistently affect the metabolic profiles of the different accessions, we used feature-based molecular networking (FBMN) to cluster compounds based on structural similarities using MS2 spectra. In this approach, networks are composed of nodes and edges. The nodes, represented by geometric shapes, symbolize each feature detected in the study. Edges are the lines that connect the nodes and represent spectral similarities between the two linked features. Each node can be further represented by pie charts, in which each slice can be colored based on the spectral count of the corresponded feature detected in different treatments or sample groups (Figure 6).

**Figure 6.**
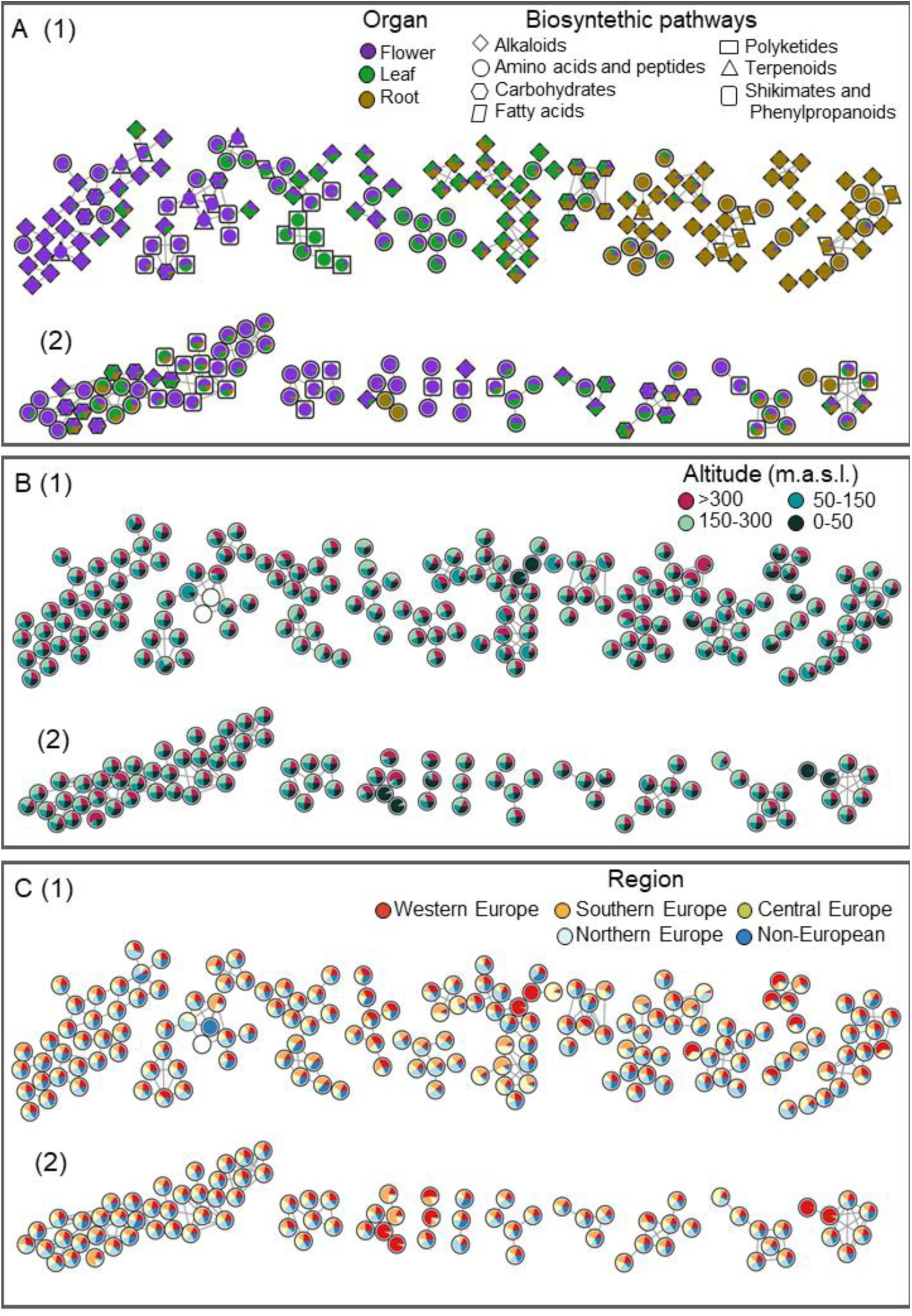
Molecular correlation networks of *Solanum dulcamara* reveal organ- and class-structured modules. Network layouts illustrate correlations among annotated features detected in (1) positive and (2) negative modes. (A) Nodes are colored by organ: flower (purple), leaf (green), root (brown), while metabolic pathways are represented by shapes. (B) Nodes are colored by altitude in meters above sea level: 0–50 (black), 50–150 (blue), 150–300 (green), >300 (red). (C) Nodes are colored by geographic region (Western, Southern, Central, Northern Europe, and non-European). Edges indicate feature-feature associations.

The FBMN of the full metabolic profiles from the accessions was used to visualize metabolic differences by organ type (flower, leaf, root), altitude of the original collected site (0-50, 50-150, 150-300, >300 m.a.s.l.), and geographical region of origin (Western Europe, Southern Europe, Central Europe, Northern Europe, and Non-European) obtained in ESI+ and ESI-datasets from all the accessions with known location of collection (Figure 6; complete networks are depicted in Supplementary Information Figure 12).

We identified organ-specific nodes across both ESI+ and ESI− datasets (Figure 6A), confirming the presence of unique features from different pathways (nodes and networks filled with only one color), as well as shared features (nodes sharing two or three colors representing the organs). However, the networks did not reveal specific metabolic differences, neither for altitude nor for regions of origin, which is observed in the majority of the nodes divided and colored among the four altitudes or the five regions assigned (Figure 6 B, C). This indicates that there is no obvious altitudinal or geographic metabolic pattern in *S. dulcamara* at the spatial scales and sampling depth that were used for this analysis.

## DISCUSSION

This study employed an untargeted LC-MS-based metabolomic approach to comprehensively investigate the chemical diversity of the wild plant species *S. dulcamara*. To characterize the plant intraspecific metabolic diversity across vegetative and reproductive organs, we grew 42 *S. dulcamara* accessions originating from different geographic locations, encompassing a wide range of altitudes, ecosystems, and climatic conditions.

### Organ-specific metabolic profiles

Our results strongly support the first hypothesis, revealing clear organ-specific metabolic profiles in *S. dulcamara*. Interestingly, leaf and flower metabolomes showed greater similarity to each other than root metabolomes, a pattern consistent across ionization modes. On the one hand, this may be because both leaf and flower spring from the apical meristem. Their formation and differentiation are tightly regulated by specific genes to coordinate shoot organ formation (Fouracre & Poethig, 2020), which makes flower organs basically morphologically modified leaves. On the other hand, this overlap may mean that above-ground organs share functional chemical traits related to dealing with aboveground conditions (UV light, heat) and biotic interactions with leaf herbivores and florivores (Kessler & Kalske, 2018). Managing this diversity of biotic interactions and dynamic abiotic conditions requires the production of a large diversity of metabolites (Walker et al., 2022; Whitehead et al., 2021). This also aligns with our observation that, overall, flowers and leaves exhibited greater metabolic richness and metabolite abundance compared to roots.

In *Solanum* species, approximately 100 steroidal alkaloids (SGAs) have been identified to date (Zhao et al., 2021), and these compounds are widely recognized for their (cyto)toxic properties. Our results support previous reports indicating that SGAs are accumulated in multiple plant organs, including roots, stems, leaves, flowers, and fruits (Calf et al., 2018; Chiocchio et al., 2023; Ginzberg et al., 2009; Shi et al., 2019; Wang et al., 2017; Zhu et al., 2018). Interestingly, SGAs represent the predominant alkaloid class in the roots, whereas they were relatively less abundant in flowers. This distribution of SGAs may align with their defensive roles against bacteria, fungi, herbivores, and other biotic threats (Calf et al., 2019; Cárdenas et al., 2019; Delbrouck et al., 2023). These may be more abundant in soils, which contain a large diversity of microbial pathogens, plant pathogenic nematodes and herbivorous soil fauna, calling for a higher level of root defenses (van Dam, 2009a). It is important to highlight that the roots analyzed in this study were adventitious roots. Previous studies have shown that SGAs profiles and concentrations differ between adventitious roots and primary roots, with primary roots generally containing higher SGAs richness than adventitious roots (Chiocchio et al., 2023). This may also explain why, overall, roots had the lowest metabolic richness in most accessions, in particular in negative mode.

Flowers, on the other hand, interact with pollinators, which may be deterred by high levels of alkaloids in floral tissues, such as the corolla or the pollen (Euler & Baldwin, 1996). In flowers, polyamines were the most abundant alkaloid compounds detected. These aliphatic nitrogen-containing compounds are known to be involved in several physiological processes critical for reproduction, including flower development, embryogenesis regulation, and protection against abiotic stress (D. Chen et al., 2019). In *Petunia* pollen, polyamines occur predominantly as hydroxycinnamoyl conjugates (phenolamides) derived from spermidine and spermine, which together contribute to pollen viability and performance. In *Petunia* cultivar V26 (violet flowers), polyamine-derived phenolamides were the most abundant alkaloids and were markedly more abundant than in pollen from cultivar W115 (white flowers) (Fernandes et al., 2026). Consistently, in *S. dulcamara* featuring mainly purple flowers, polyamines represented the predominant alkaloid. Polyamines play an important role in reproduction; they are recognized as major constituents of floral pollen, with remarkable contributions to pollen function and fertility (Spremo et al., 2025).

We also found relatively high levels of shikimates and phenylpropanoids, mainly detected in ESI-mode. Across all three organs, cinnamic acid derivatives represented the most abundant phenylpropanoid class, while flavonols were particularly abundant in flowers and roots. Cinnamic acid derivatives are widely recognized for their antimicrobial and antifungal properties, which affect membrane permeability and interfere with enzymatic activity, while also contributing to plant growth and development (Shuab et al., 2016). Flavonols, a major subclass of flavonoids, are crucial for photoprotection, antioxidant responses, and developmental regulation, playing important regulatory roles during plant maturation and tissue differentiation (Mahmud et al., 2023). Interestingly, two accessions with white flowers showed an increased abundance of several flavonoids, compared to the purple flowered accessions in the set. From the well-studied model *Petunia hybrida* white flowers typically result from impaired anthocyanin accumulation (i.e., reduced production of purple pigments), leading to increased flavanones and flavonols (Pollastri & Tattini, 2011). Floral flavonoids have been linked to pollinator preferences in *Petunia* plants, with flavonols as driver compounds to attract nocturnal visitors to *Petunia axillaris* (Hoballah et al., 2005). A potential metabolic trade-off associated with flower color has been proposed in *Petunia*, and a similar trade-off may also occur in *S. dulcamara*.

Our comprehensive analysis also revealed the importance of employing both positive and negative modes when obtaining a comprehensive metabolic atlas. The differential detection efficiency is attributed to the inherent chemical properties such as electronegativity, basicity, and size of the molecule analyzed. In the compound classes we were interested in, alkaloids generally favor protonation in ESI+ and phenolic/acidic compounds favoring deprotonation in ESI-. By combining positive and negative ionization modes, we thus obtained a more comprehensive profile (Liigand et al., 2017).

### Steroidal glycoalkaloids (SGAs) as drivers of chemodiversity and chemotypes

Our results strongly supported the hypothesis that SGAs are major drivers of *S. dulcamara* chemodiversity, but only in flowers and leaves. Based on SGA data, leaf and flower chemical profiles formed four clear clusters, or chemotypes (Müller et al., 2026) aligning with previously described *S. dulcamara* chemotypes. In both leaves and flowers, the SGAs profiles revealed distinct chemotypes, as reported by Calf et al. (2018), who sampled accessions originating from populations in the Netherlands. Interestingly, expanding our collection towards other accessions from very different locations did not increase the number of previously described leaf chemotypes, consistent mainly in accessions with SGAs whose aglycones are predominantly saturated or unsaturated (Calf et al., 2018, 2019; Willuhn, 1966). Apart from two accessions originating from a single Dutch population which lacked SGAs, all accessions grouped into the other three distinct SGA leaf and flower chemotypes, based on the structure of the steroidal aglycon. Additional ecologically relevant chemical differentiation in SGA profiles is generated by adding different types and numbers of sugar moieties to the aglycon (Itkin et al., 2013). In potato it was found that the transition from a triose (three sugars) to a tetraose (four sugars) can make potato leaves resistant to Colorado potato beetles and a microbial pathogen (Wolters et al., 2023). Based on the high-resolution mass spectrometric data and retention time information, as well as the fragmentation patterns and sugar losses obtained in our tandem MS/MS analysis, we could also annotate both trioses and tetraoses among the putatively annotated SGAs, which were present in different aglycones and isomers.

Our analyses showed that accessions ZD11 and ZD04 were separated from the remaining accessions due to having very low levels of SGAs. This partly contrasts with earlier reports, in which it was reported that ZD04 contains SGAs (Calf et al., 2018, 2019). The individuals from those accessions have been maintained, propagated, and moved for almost a decade, increasing the possibility of label mix-ups. Alternatively, long-term cultivation under controlled conditions could cause metabolic adaptation over time, resulting in shifts in their chemical profiles.

In contrast, root SGAs profiles were rather uniform and did not separate into particular chemotypes except for the accessions ZD11 and ZD04. Chemical profile differences among tissues or organs are not uncommon; other studies have demonstrated that chemical diversity varies among above- and belowground tissues as a result of different selection pressures (Kleine & Müller, 2013; Müller et al., 2026; Rahimova et al., 2025). Similarly, no chemotypes clustering was observed in the flavonoids and cinnamic acid derivatives profiles (Calf et al., 2018, 2019)

### Influence of Geographic Origin and Environmental Factors

Contrary to our hypothesis, and using a molecular network approach, this study found no consistent metabolomic variation associated with the geographic origin or altitude among the *S. dulcamara* accession we analyzed but confirmed that there are metabolite clusters associated with plant organs. This hypothesis was formulated considering that significant differences in the metabolic profile are commonly observed in studies analyzing accessions from different geographical origins (Lin et al., 2023; Skubel et al., 2020) or altitudes (Nomoto et al., 2025). However, in our study, neither the geolocation nor the altitude related to the geographical origin of the seeds were associated with any specific metabolic signature. Although the sample set in the present study spans a wide latitudinal, longitudinal and altitudinal range across the Northern Hemisphere, the exact geolocations and altitudinal origin were not always known to detail. Assessing the effects of geographical location and altitude may require a more extensive and targeted sampling effort. In general, it should include the analysis of multiple individuals from several populations collected at different locations, which was beyond the scope of this study.

## CONCLUSIONS

Using LC-MS-based metabolic data of roots, leaves and flowers, we characterized the metabolic diversity of different *S. dulcamara* organs. Thereby, we found different levels of intraspecific chemodiversity on the organ and accession level. Overall, we could assign the accessions to four main leaf and flower SGA chemotypes, whereas the accessions did not cluster in clear chemotypes based on flavonoids or cinnamic acid derivatives profiles in any organ. Our data are the basis for unraveling possible functional roles of such chemodiversity in mediating interactions between plants, environmental conditions and other organisms. As a wild relative of cultivated Solanaceae species, including tomato, potato, and eggplant, the chemical characterization of *S. dulcamara* may harbor metabolites with agronomic relevance, such as resistance factors and (anti)nutritional traits. In addition to its ecological relevance, therefore, this study may contribute to applied research in plant breeding and crop production.

## SUPPLEMENTARY INFORMATION

Supplementary Figure 1. Hierarchical cluster analysis of *Solanum dulcamara* metabolomics data acquired in ESI+.

Supplementary Figure 2. Hierarchical cluster analysis of *Solanum dulcamara* metabolomics data acquired in ESI-.

Supplementary Figures 3-11. Organ-wise clustering of major specialized metabolite classes across *Solanum dulcamara* genotypes.

Supplementary Figure 12. Full feature based molecular networks (FBMN) obtained by GNPS.

Supplementary Information S1. Metadata

Supplementary Information S2. POS_NPC#Pathway

Supplementary Information S3. NEG_NPC#Pathway

Supplementary Information S4. POS_Richness and Abundance

Supplementary Information S5. NEG_Richness and Abundance

Supplementary Information S6. SGAs_list

Supplementary Information S7. Flavonoids_list

Supplementary Information S8. Cinnamic acid derivatives_list

## AUTHOR CONTRIBUTIONS

NMvD initiated the study. JVMS, PCPB and NMvD designed the study. JVMS prepared and collected the plants, JVMS and PCPB conducted phytochemical analysis, JVMS, PCPB and AMP analyzed LC-MS data and prepared figures. JVMS, AMP, PCPB and NMvD wrote and agreed on the final manuscript.

## DATA AVAILABILITY

Data are available in the Supplementary information. All relevant raw data sets are available upon request. The GNPS Jobs and parameters can be found at Task IDs: caf1969ae09e4d18b3883744ce2cc3e0 and 99ddf56bf20b4347a15a397cc6e5f10d.

## ACKNOWLEDGEMENTS

We thank Dr. Khabat Vahabi from the SSP group in IGZ for providing guidance on statistical approaches and data processing and Prof. Dr. Anke Steppuhn and MSc. Kruthika Sen Aragam from the University of Hohenheim for sharing accessions from their collection. Finally, we thank Andrea Janskowsky (SSP) and the gardeners at IGZ for expert support.

