## Supplementary_Information_Figures for "Untargeted metabolic analysis reveals intraspecific and organ-specific chemodiversity in *Solanum dulcamara*"

### Equal last authors

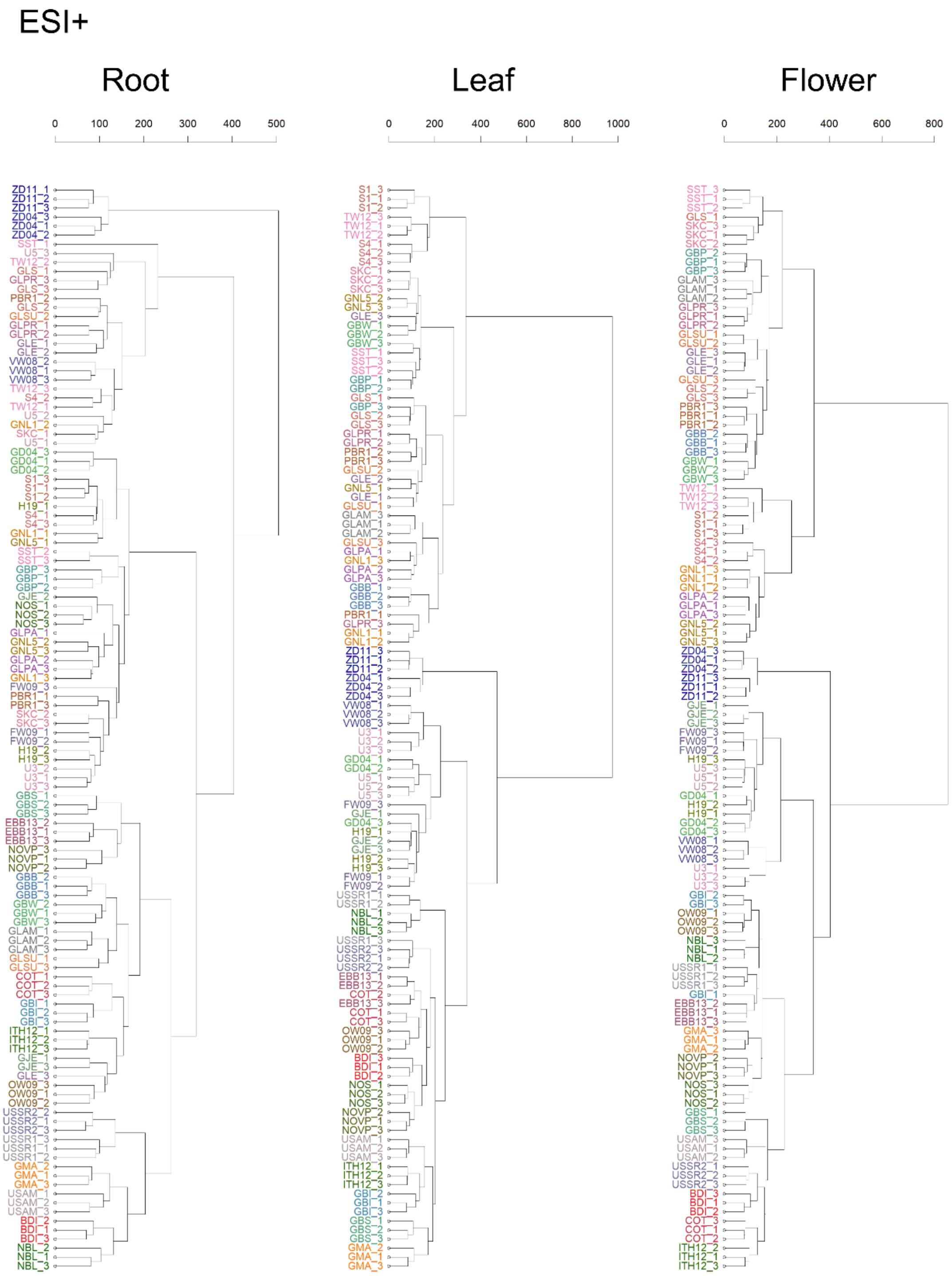


**Supplementary Figure 1. Hierarchical cluster analysis of *Solanum dulcamara metabolomics data* acquired in ESI+**. Euclidean distance was used as the similarity measure, and clusters were generated using Ward's linkage method.

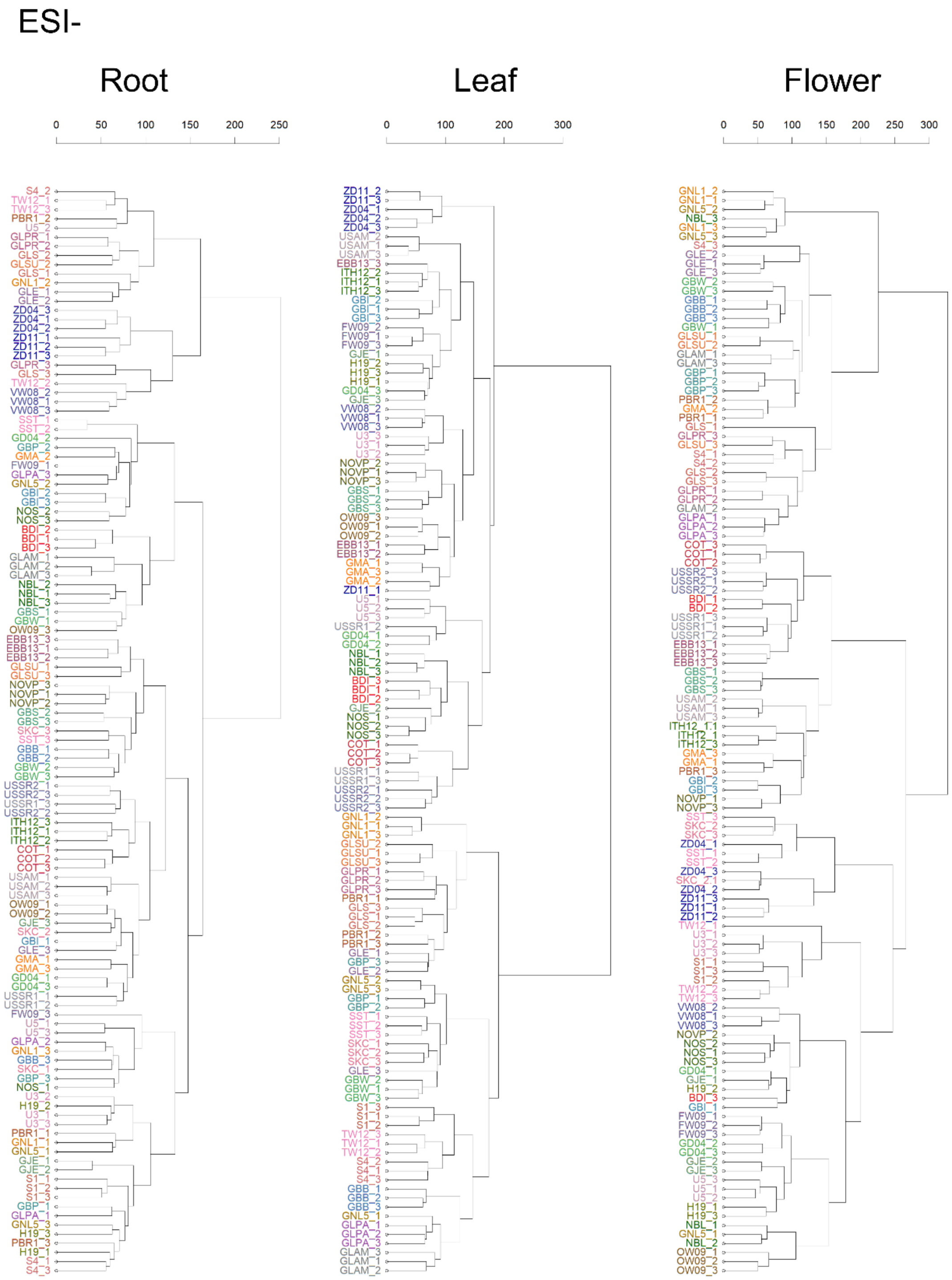


**Supplementary Figure 2. Hierarchical cluster analysis of *Solanum dulcamara* metabolomics data acquired in ESI-**. Euclidean distance was used as the similarity measure, and clusters were generated using Ward's linkage method.

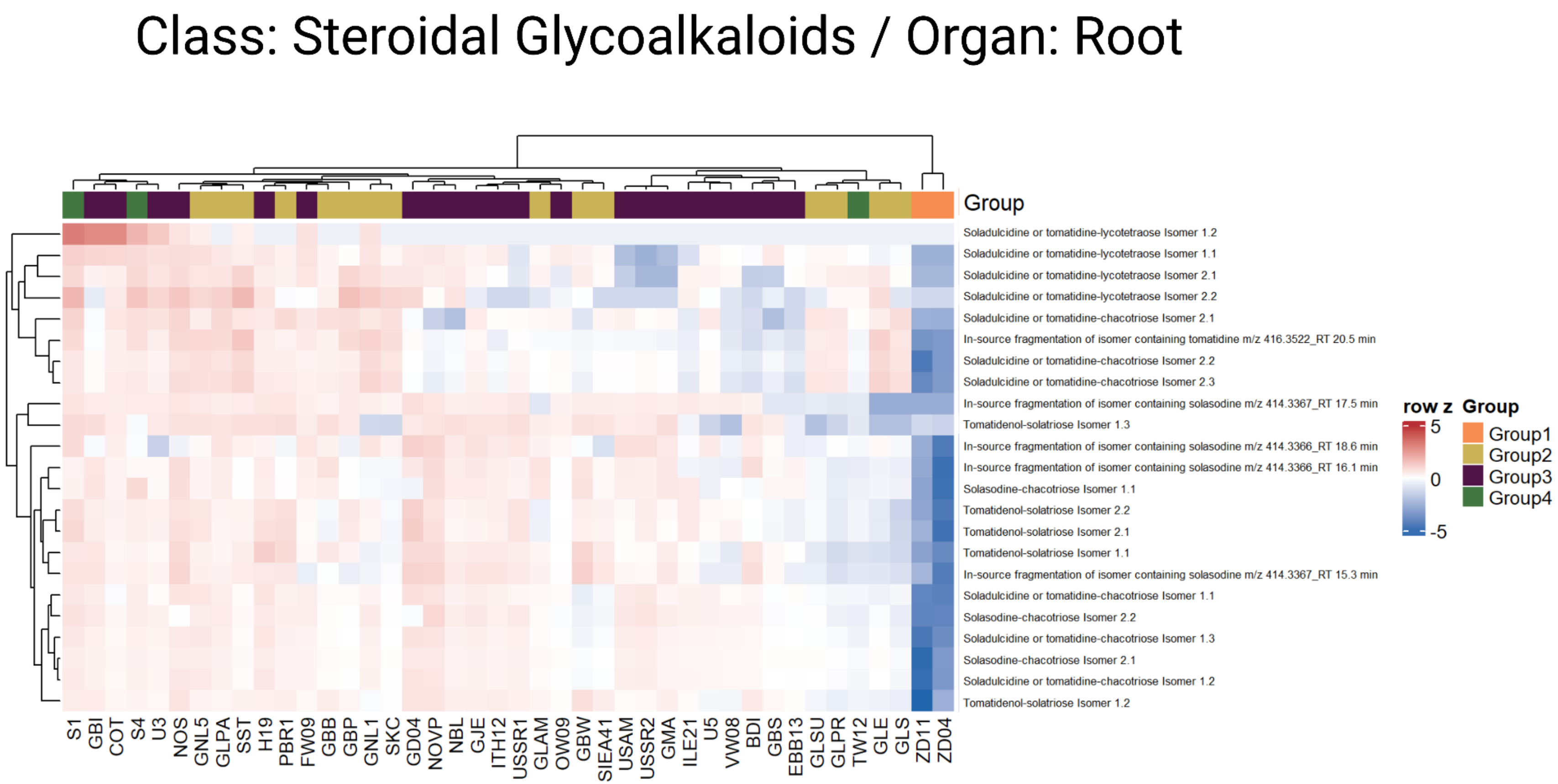


**Supplementary Figure 3. Organ-wise clustering of major specialized metabolite classes across *Solanum dulcamara* genotypes.** Hierarchically clustered heatmaps show standardized metabolite levels (row z-scores; blue = lower, red = higher) for steroidal glycoalkaloids (SGAs) in roots. Rows represent metabolites and columns represent genotypes. Dendrograms indicate similarity-based clustering, while the upper bars denote genotype groups (Group 1–4).

### Class: Steroidal Glycoalkaloids / Organ: Leaf

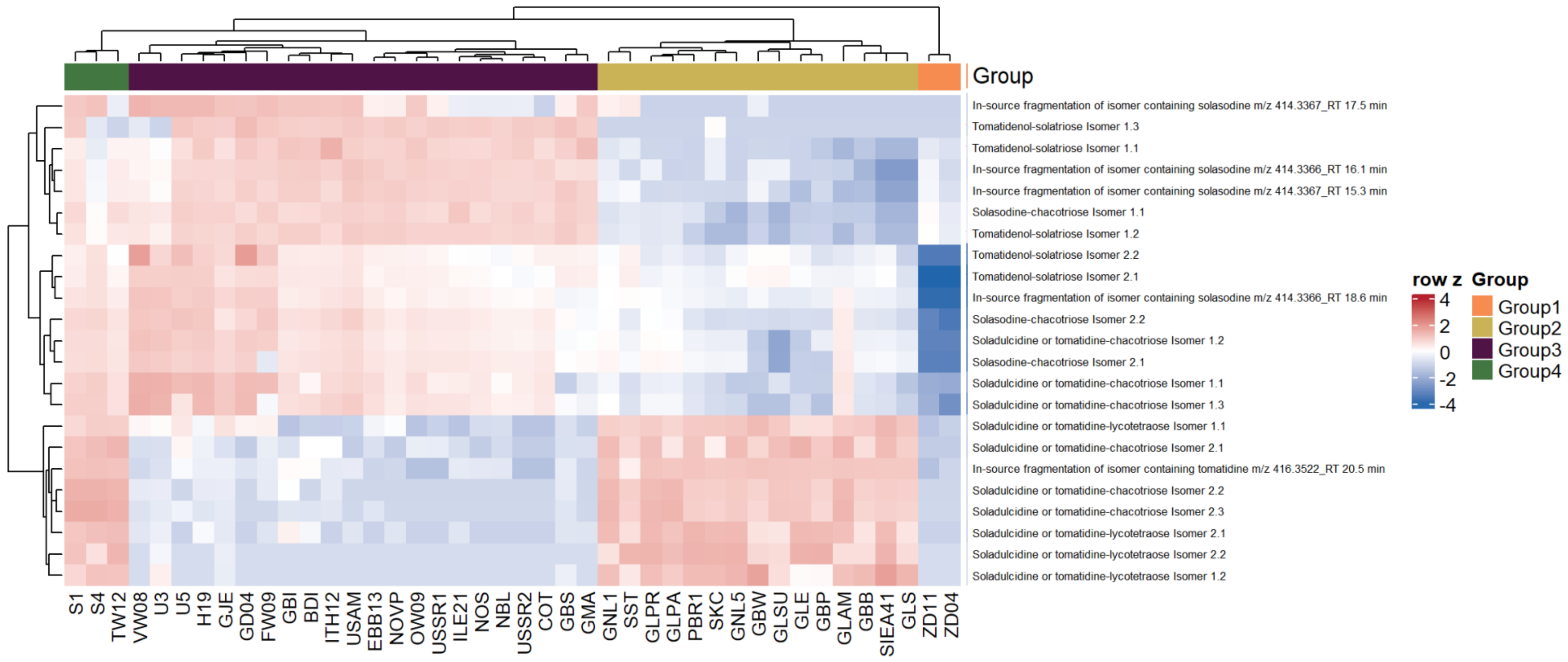


**Supplementary Figure 4. Organ-wise clustering of major specialized metabolite classes across *Solanum dulcamara* genotypes.** Hierarchically clustered heatmaps show standardized metabolite levels (row z-scores; blue = lower, red = higher) for steroidal glycoalkaloids (SGAs) in leaves. Rows represent metabolites and columns represent genotypes. Dendrograms indicate similarity-based clustering, while the upper bars denote genotype groups (Group 1–4).

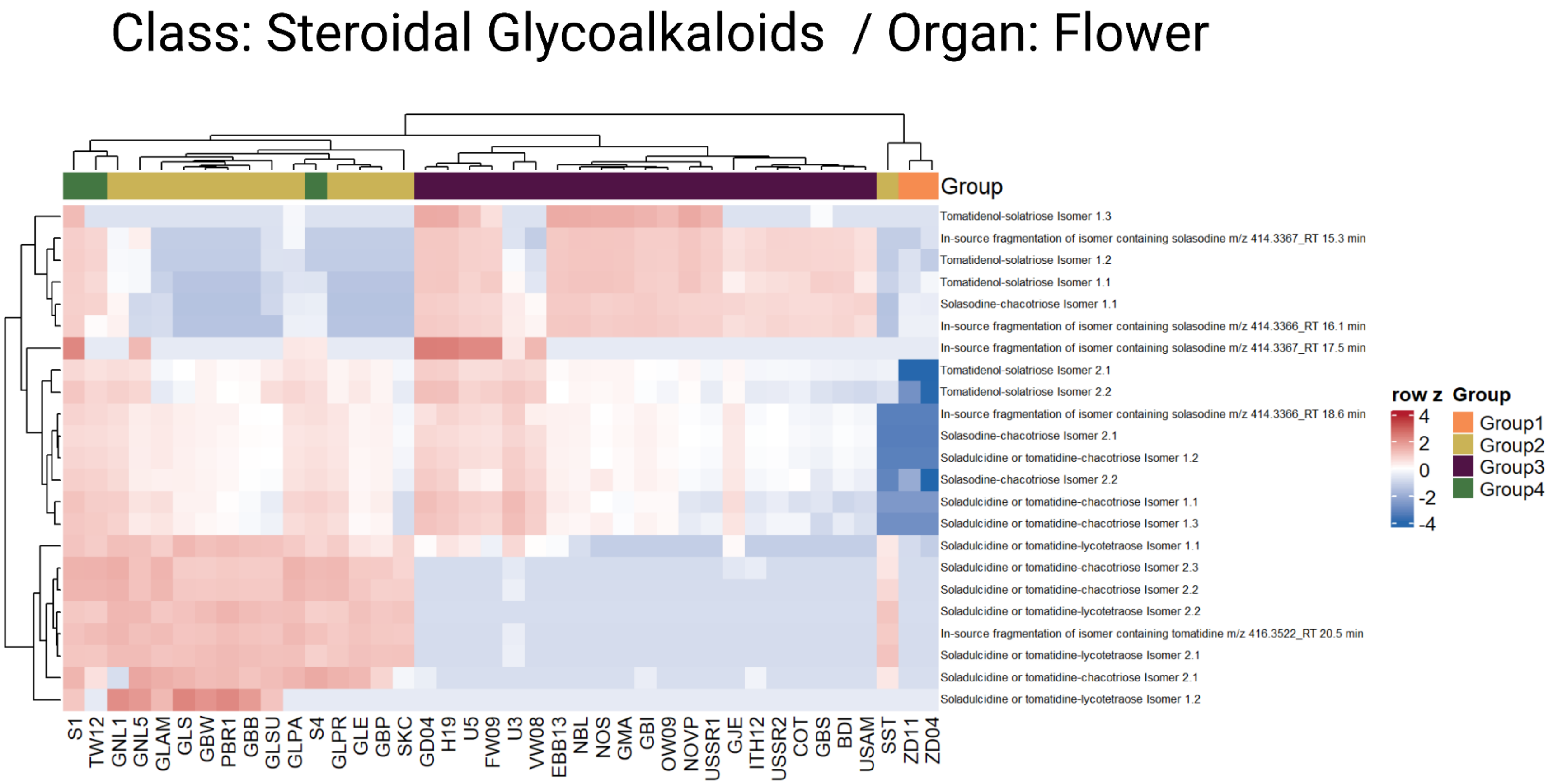


**Supplementary Figure 5. Organ-wise clustering of major specialized metabolite classes across *Solanum dulcamara* genotypes.** Hierarchically clustered heatmaps show standardized metabolite levels (row z-scores; blue = lower, red = higher) for steroidal glycoalkaloids (SGAs) in flowers. Rows represent metabolites and columns represent genotypes. Dendrograms indicate similarity-based clustering, while the upper bars denote genotype groups (Group 1–4).

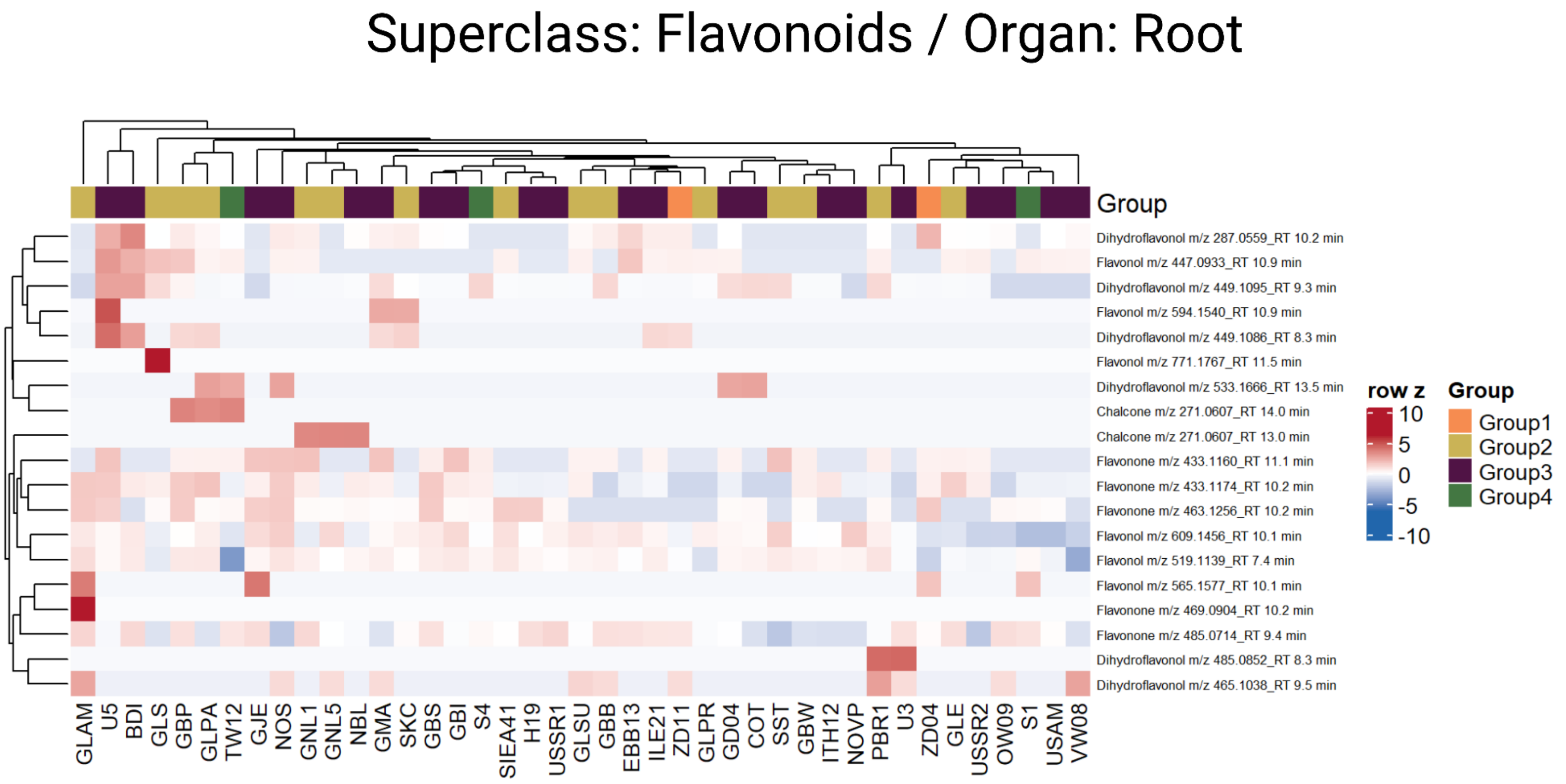


**Supplementary Figure 6. Organ-wise clustering of major specialized metabolite classes across *Solanum dulcamara* genotypes.** Hierarchically clustered heatmaps show standardized metabolite levels (row z-scores; blue = lower, red = higher) for flavonoids in roots. Rows represent metabolites and columns represent genotypes. Dendrograms indicate similarity-based clustering, while upper bars denote genotype groups (Group 1–4).

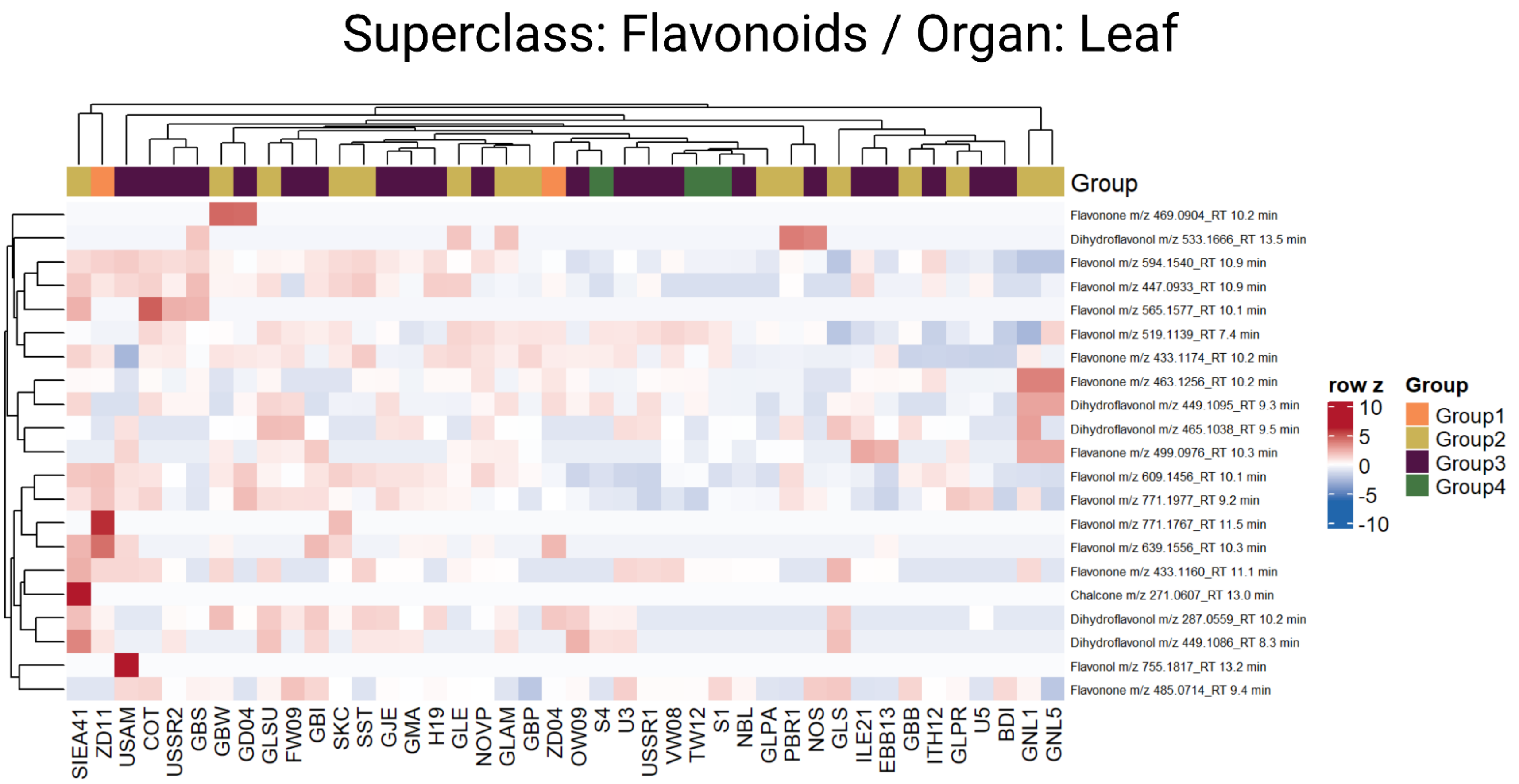


**Supplementary Figure 7. Organ-wise clustering of major specialized metabolite classes across *Solanum dulcamara* genotypes.** Hierarchically clustered heatmaps show standardized metabolite levels (row z-scores; blue = lower, red = higher) for flavonoids in leaves. Rows represent metabolites and columns represent genotypes. Dendrograms indicate similarity-based clustering, and upper bars denote genotype groups (Group 1–4).

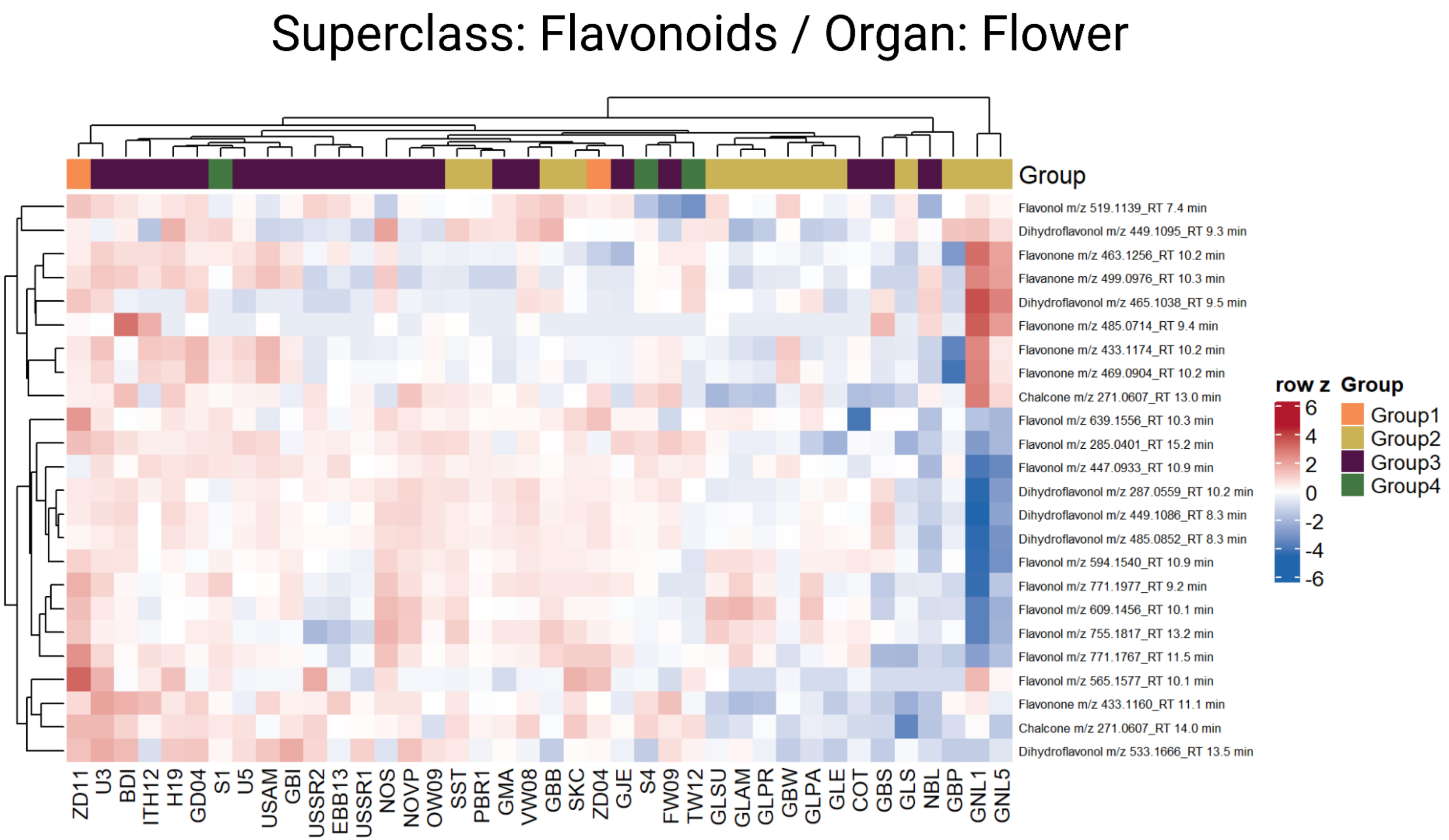


**Supplementary Figure 8. Organ-wise clustering of major specialized metabolite classes across *Solanum dulcamara* genotypes.** Hierarchically clustered heatmaps show standardized metabolite levels (row z-scores; blue = lower, red = higher) for flavonoids in flowers. Rows represent metabolites and columns represent genotypes. Dendrograms indicate similarity-based clustering, and upper bars denote genotype groups (Group 1–4).

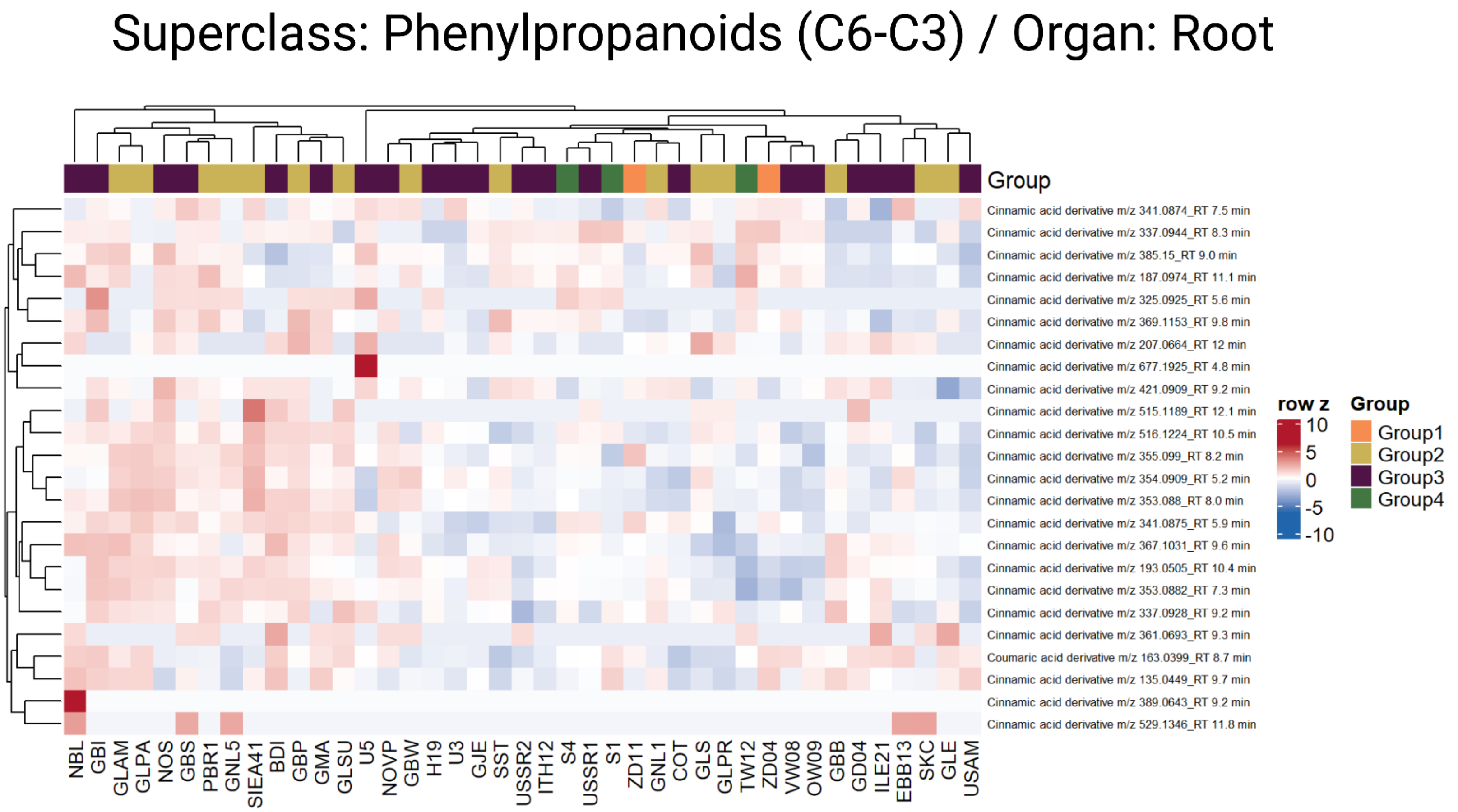


**Supplementary Figure 9. Organ-wise clustering of major specialized metabolite classes across *Solanum dulcamara* genotypes.** Hierarchically clustered heatmaps show standardized metabolite levels (row z-scores; blue = lower, red = higher) for phenylpropanoids (C6-C3) in roots. Rows represent metabolites and columns represent genotypes; dendrograms indicate similarity-based clustering, and upper bars denote genotype groups (Group 1–4).

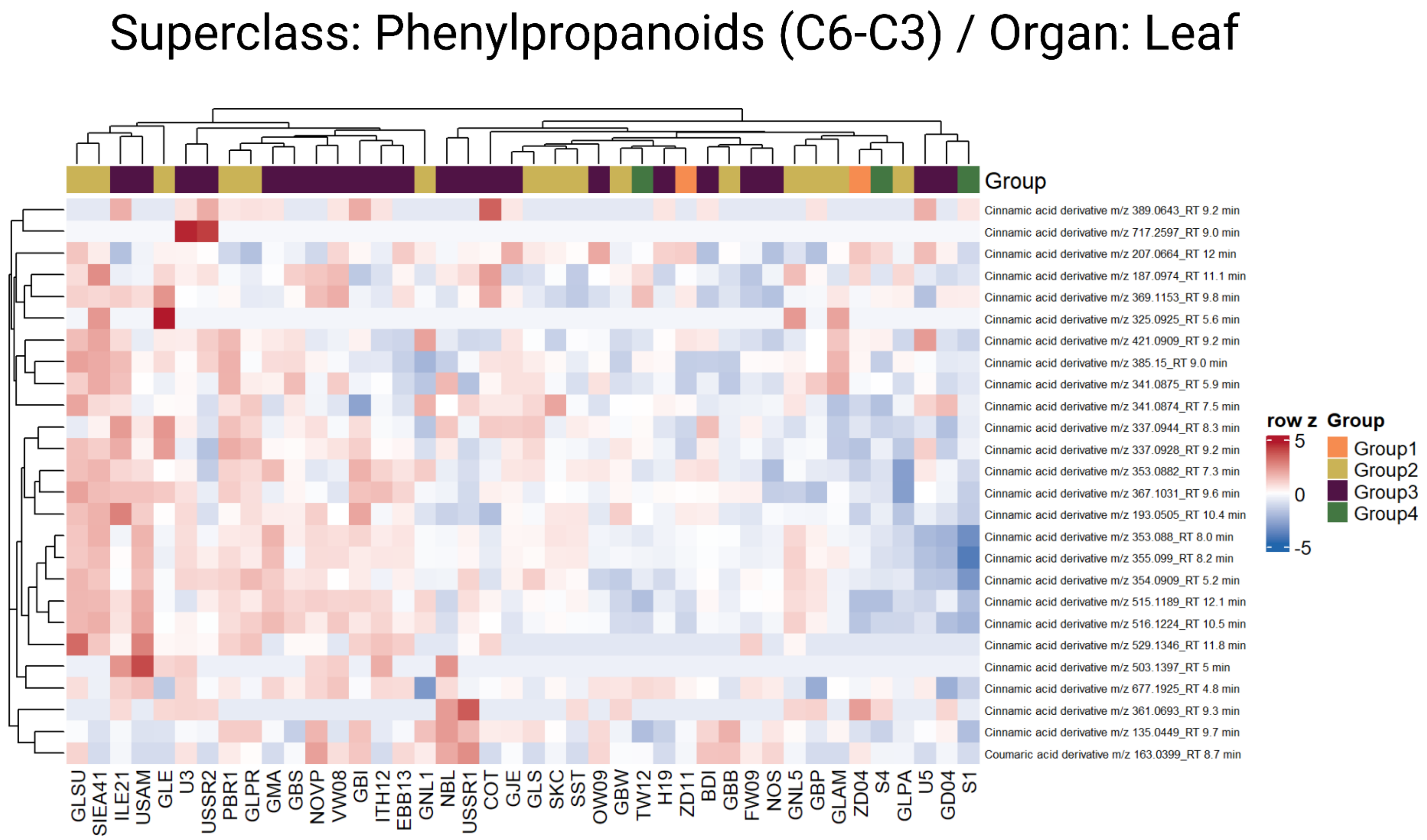


**Supplementary Figure 10. Organ-wise clustering of major specialized metabolite classes across *Solanum dulcamara* genotypes.** Hierarchically clustered heatmaps show standardized metabolite levels (row z-scores; blue = lower, red = higher) for phenylpropanoids (C6-C3) in leaves. Rows represent metabolites and columns represent genotypes; dendrograms indicate similarity-based clustering, and upper bars denote genotype groups (Group 1–4).

### Superclass: Phenylpropanoids (C6-C3) / Organ: Flower

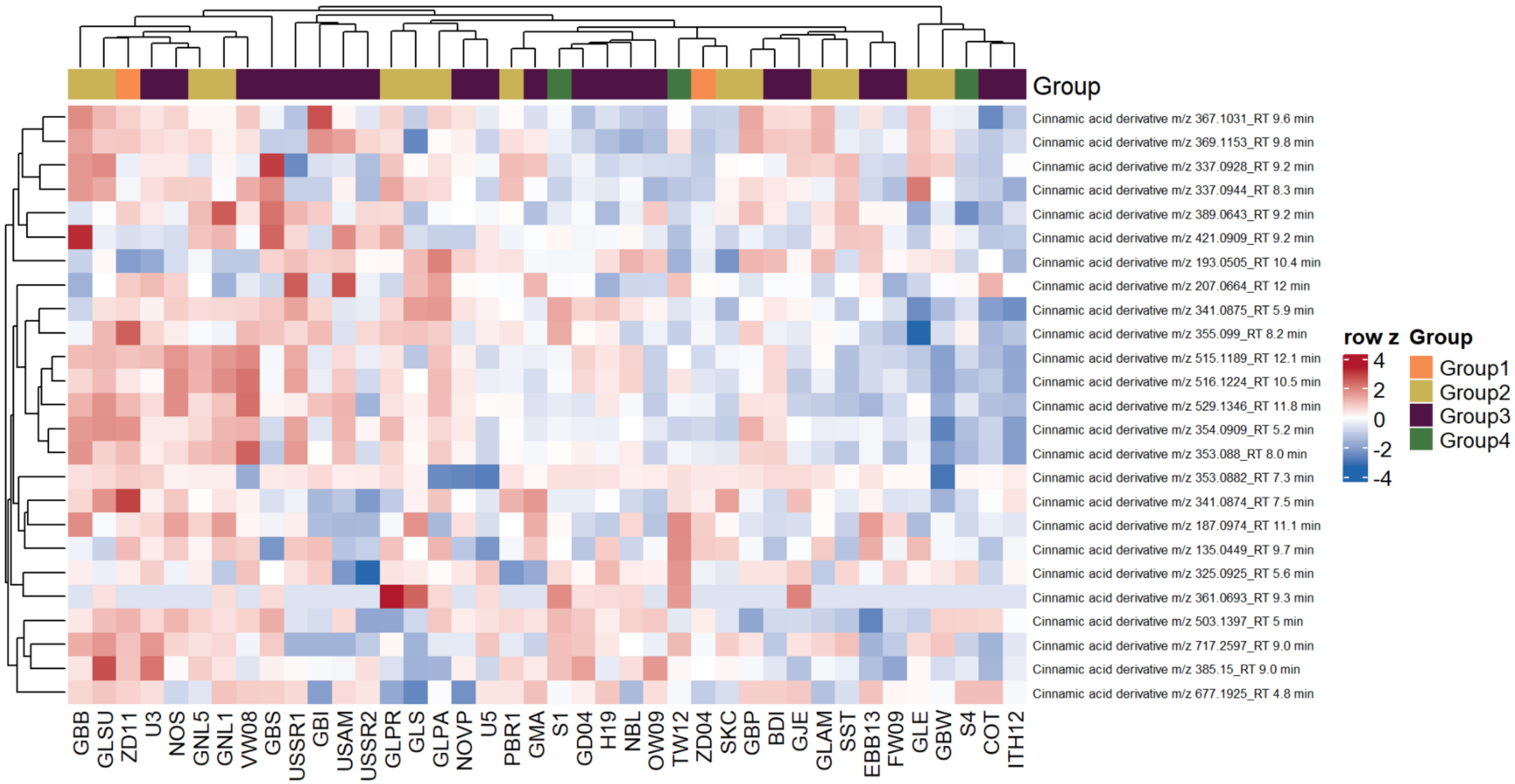


**Supplementary Figure 11. Organ-wise clustering of major specialized metabolite classes across *Solanum dulcamara* genotypes.** Hierarchically clustered heatmaps show standardized metabolite levels (row z-scores; blue = lower, red = higher) for phenylpropanoids (C6-C3) in flowers. Rows represent metabolites and columns represent genotypes; dendrograms indicate similarity-based clustering, and upper bars denote genotype groups (Group 1–4).

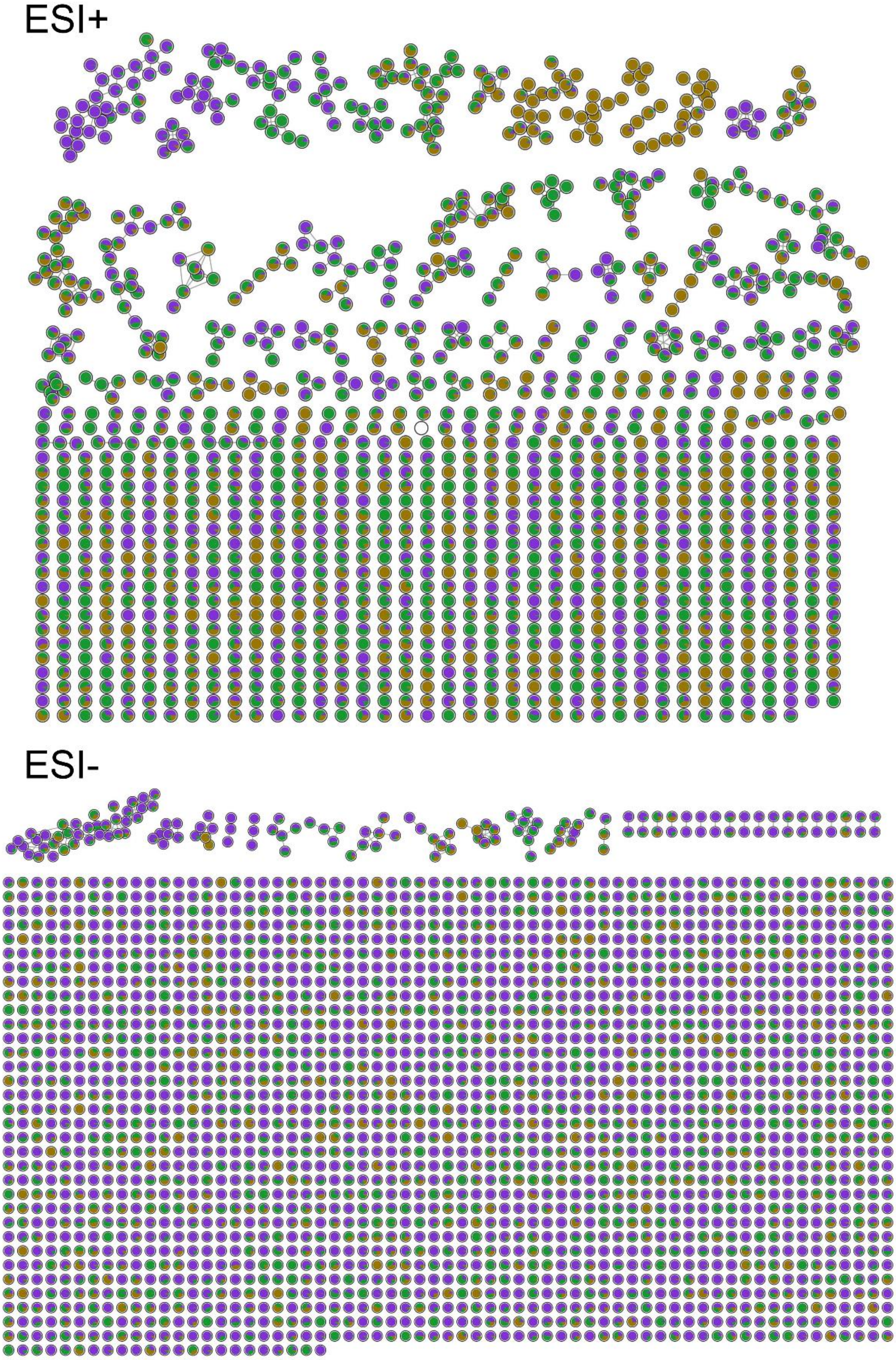


**Supplementary Figure 12. Full feature-based molecular networks (FBMN) of *Solanum dulcamara* obtained by GNPS.** Network layouts with nodes colored by organ (flower, leaf, root). ESI+: FBMN obtained in positive mode; ESI-: FBMN obtained in negative mode.
